# An innovative hematopoietic stem cell gene therapy approach benefits CLN1 disease in the mouse model

**DOI:** 10.1101/2022.03.03.482460

**Authors:** Marco Peviani, Sabyasachi Das, Janki Patel, Odella Jno-Charles, Rajesh Kumar, Ana Zguro, Tyler D. Mathews, Rita Milazzo, Eleonora Cavalca, Valentina Poletti, Alessandra Biffi

## Abstract

Hematopoietic stem and progenitor cells (HSPCs) can lead to the establishment of a long-lasting microglia-like progeny in the brain of properly myeloablated hosts. We exploited this approach to treat the severe CLN1 neurodegenerative disorder, which is the most aggressive form of neuronal ceroid lipofuscinoses, due to deficiency of palmitoyl-protein thioesterase 1 (hPPT1). We here provide first evidence that: i) transplantation of wild type HSPCs exerts a partial but long-lasting mitigation of the symptoms; ii) transplantation of HSPCs over-expressing hPPT1 by lentiviral gene transfer enhances therapeutic benefit as compared to wild type cell transplant, with first demonstration of such a dose-effect benefit for a purely neurodegenerative condition like CLN1 disease; iii) transplantation of hPPT1 over-expressing HSPCs by a novel intracerebroventricular (ICV) approach is sufficient to transiently ameliorate CLN1 disease symptomatology in the absence of hematopoietic tissue engraftment of the transduced cells; and iv) the combinatorial transplantation of transduced HSPCs intravenously and ICV results in the most robust therapeutic benefit among the tested approaches on both pre-symptomatic as well as symptomatic animals. Overall, these findings provide first evidence of the efficacy and feasibility of this novel approach to treat CLN1 disease and possibly other neurodegenerative conditions, paving the way for its future clinical application.

## Introduction

Neuronal Ceroid Lipofuscinoses (NCLs or CLNs) are a group of genetic disorders characterized by neurodegeneration and intracellular accumulation of an auto-fluorescent lipopigment^1^. Together, they represent the most prevalent class of childhood neurodegenerative diseases. The NCLs encompass several distinct biological entities that vary in age at onset, specific neurological phenotype and rate of progression. These entities share characteristic clinical features such as progressive vision loss, dementia, epileptic seizures and loss of motor coordination, culminating in premature death. CLN1 disease, first described and one of the most frequent causes of NCL, is a lysosomal storage disorder (LSD) inherited in an autosomal recessive manner^2,3^. It is due to biallelic loss-of-function variants in the *PPT1* gene (also designated CLN1; 1p32) encoding the lysosomal enzyme palmitoyl-protein thioesterase 1 (PPT1) that cleaves the thioester linkage of the fatty acid palmitate to cysteine residues of palmitoylated proteins (constituents of ceroid), a step that is necessary to allow degradation of these proteins by lysosomes. In most of the cases, CLN1 disease manifests clinically as infantile NCL (INCL) that is characterized by onset during childhood, at around 6-24 months of age, and has an invariably fatal outcome by 9-13 years of age^4^. However, juvenile-onset and adult-onset forms of the disease have been described. Currently, no curative treatment is available that could prevent or reverse this fatal course.

The development of treatments for infantile CLN1 disease has been greatly accelerated by the availability of the Ppt1^-/-^ mouse model, where the murine gene coding for *Ppt1* has been knocked-out^5,6^. Ppt1^-/-^ mice recapitulate closely many phenotypic features observed in patients, including progressive motor disability^5^, visual impairment^7^, muscle twitches and recurrent seizures^5,8^. At a histopathological level, they accumulate autofluorescent storage (AF) material in neurons and glial cells, show progressive neuronal loss (leading to thinning of the cortical layer) and prominent neuroinflammation (astrogliosis and microgliosis)^5,9–12^.

Several therapeutic strategies have been tested in Ppt1^-/-^ mice with variable outcomes, including symptomatic treatments addressing the clinical manifestations of the disease, such as anti-excitotoxicity drugs;^8^ substrate reduction therapy exploiting cysteamine bis-tartrate or N-tert-butyl hydroxylamine^13,14^; or enzyme replacement therapy.^15^ In the latter case, insufficient widespread reconstitution of PPT1 activity in the CNS, including brain and spinal cord, is likely the key limitation that hindered successful clinical application by now. One major breakthrough has been recently reported at preclinical level by exploiting a gene therapy approach based on adeno-associated viral vectors (AAV) expressing the wild type PPT1 administered in the cerebrospinal fluid in Ppt1^-/-^ mice.^16^ However, as per most recent indications for the development of novel treatment options for LSDs, targeting secondary disease mechanisms, such as microglia activation and neuroinflammation, in addition to intervening on the primary disease culprit by compensating the enzymatic deficiency, is warranted for achieving robust therapeutic benefit^12,17,18^. In this setting, microglia cells, which are strongly implicated in the pathogenesis of LSDs, should be considered as a primary target cell type.

Hematopoietic stem cell (HSC) gene therapy may successfully address these indications. Indeed, transplantation of genetically engineered hematopoietic stem and progenitor cells (HSPCs) in appropriate conditions can generate a microglia-like progeny that restores an efficient scavenging function within the whole CNS, with the potential to clear some of the accumulated substrate, reverse the detrimental effects of microglia activation due to the accumulation of un-degraded substrate and provide a new effective source of bioavailable enzyme within the CNS^19^. Here, we provide the first proof of applicability and therapeutic efficacy of HSC gene therapy in the CLN1 disease mouse model and exploit a newly developed CNS-targeting modality to further enhance HSC therapeutic effects.

## Results

### A novel disease severity scoring system accurately measures CLN1 disease symptoms in Ppt1^-/-^ mice

Identification of functional and clinically relevant pathologic readouts is instrumental to assess the efficacy of new therapeutic approaches. The motor function and behavioral assessment that includes the rotarod and/or the open field tests^5,10,15^ to describe the Ppt1^-/-^ mice phenotype highlights significant deterioration of motor performance starting from around 210 days of age. Afterwards, a very steep worsening of symptoms is observed, followed by a stable poor performance up until the end stage of disease, with death occurring at around 245-260 days of age. At close examination, Ppt1^-/-^ mice display early symptoms starting from 140-150 days of age, including pop-corn seizures and jerks, abnormal displacement of hind-and/or fore-limbs when raised by the tail, alteration of the grooming behavior, that are not measured by the rotarod and open field tests. We therefore combined all these observations into a scoring system (see Fig. S1A and supplementary table 1) that provides a more comprehensive characterization of the phenotype of Ppt1^-/-^ mice from the early stages of disease progression. Given that the score increases with worsening of symptoms, we called it a disease severity score (DSS).

As shown in Fig. S1B, wild type (WT) mice never display DSS values higher than 1, whereas Ppt1^-/-^ mice show a DSS ∼2 starting at ∼160 days of age. A significant worsening of Ppt1^-/-^ mice symptoms then happens and by 190 days of age animals display a rapid increase of DSS, followed by a progressive deterioration of the overall body condition up to 225 days, when they begin to reach the humane end point (HEP) (DSS ∼6-7 on average). Rotarod testing conducted in parallel to DSS evaluation (Fig. S1C) confirmed that the latter detects disease manifestations earlier than the former. Overall, based on DSS we could identify three disease stages, characterized by specific behavioral and motor deficits, associated with the worsening of symptoms as the disease progresses (Fig. S1D): stage 1 or onset, defined by a DSS ≥2 at median age of 173 days; stage 2 or symptomatic phase, defined by a DSS ≥4 at median age of 205 days; stage 3 or end stage, with a DSS ≥6 at median age of 229 days. We used this classification system to evaluate the effects of HSPC transplantation and HSC gene therapy in Ppt1^-/-^ mice.

### Transplantation of WT hematopoietic cells attenuates the phenotype of Ppt1^-/-^ mice

To generate early feasibility data on HSPC transplantation in CLN1 disease, 6-8 weeks-old Ppt1^-/-^ mice were exposed to a busulfan-based myeloablative conditioning and then transplanted intravenously (IV) with either unmanipulated total bone marrow (BM) or lineage negative (Lin^-^) HSPCs, corresponding to the CD34^+^ fraction in humans, retrieved from WT Ppt1^+/+^ donors (Fig. S2A). To track donor-derived cells *in vivo*, Lin^-^ HSPCs were transduced with a lentiviral vector (LV) expressing the D.NGFR (Nerve Growth Factor Receptor) reporter gene, whereas for tracking BM cells we relied on an allelic mismatch at the CD45 locus between donors (CD45.1) and recipients (CD45.2). As control groups, Ppt1^-/-^ mice were left untreated (UT) or were mock transplanted (namely, underwent conditioning and transplantation of Ppt1^-/-^ un-manipulated, un-transduced BM), to monitor the effects of conditioning on the disease. Starting at 140 days of age (85-100 days post-transplant) animals were assessed weekly for DSS, to monitor disease progression. Mock-transplanted animals displayed an earlier symptom onset, with DSS >2 already at 154 days of age (Fig. 1A) and higher thereafter, and a shorter survival (median survival 228 versus 248 days, respectively; p <0.0001 Log-rank) compared to UT controls (Fig. 1B). This could be due to the well-known intrinsic toxicity of the conditioning *per se* and of the pharmacological-grade busulfan formulation used for conditioning (Busilvex™, Pierre Fabre Médicament), that contains dimethyl-acetamide^23^.Ppt1^-/-^ mice transplanted with WT BM or Lin^-^ cells displayed a milder disease progression than mock-transplanted and UT controls. Treated animals had DSS values below 4 up until 224 days of age (study termination, when the first UT transplanted control died, dashed line in Fig. 1B) and a significantly ameliorated DSS trajectory over time (Fig. 1A). Disease progression was superimposable between Ppt1^-/-^ mice transplanted with Lin^-^ or BM WT cells (Fig. S2B). Given the improved behavior of the transplanted mice, a small group of animals was kept alive (n=7) for long-term observation to confirm that the disease remained mild (DSS below 6) beyond the natural disease course of Ppt1^-/-^ UT controls (of 230-237 days of age, on average). Except for two animals sacrificed early due to skin wounds (asterisk in Fig. 1B), the remaining animals survived up to at least 280 days of age, with a DSS ≤5 (Fig. S2B) and without limbs stiffness. The last animals were finally euthanized at 330 days of age without reaching the HEP.

**Fig 1.**
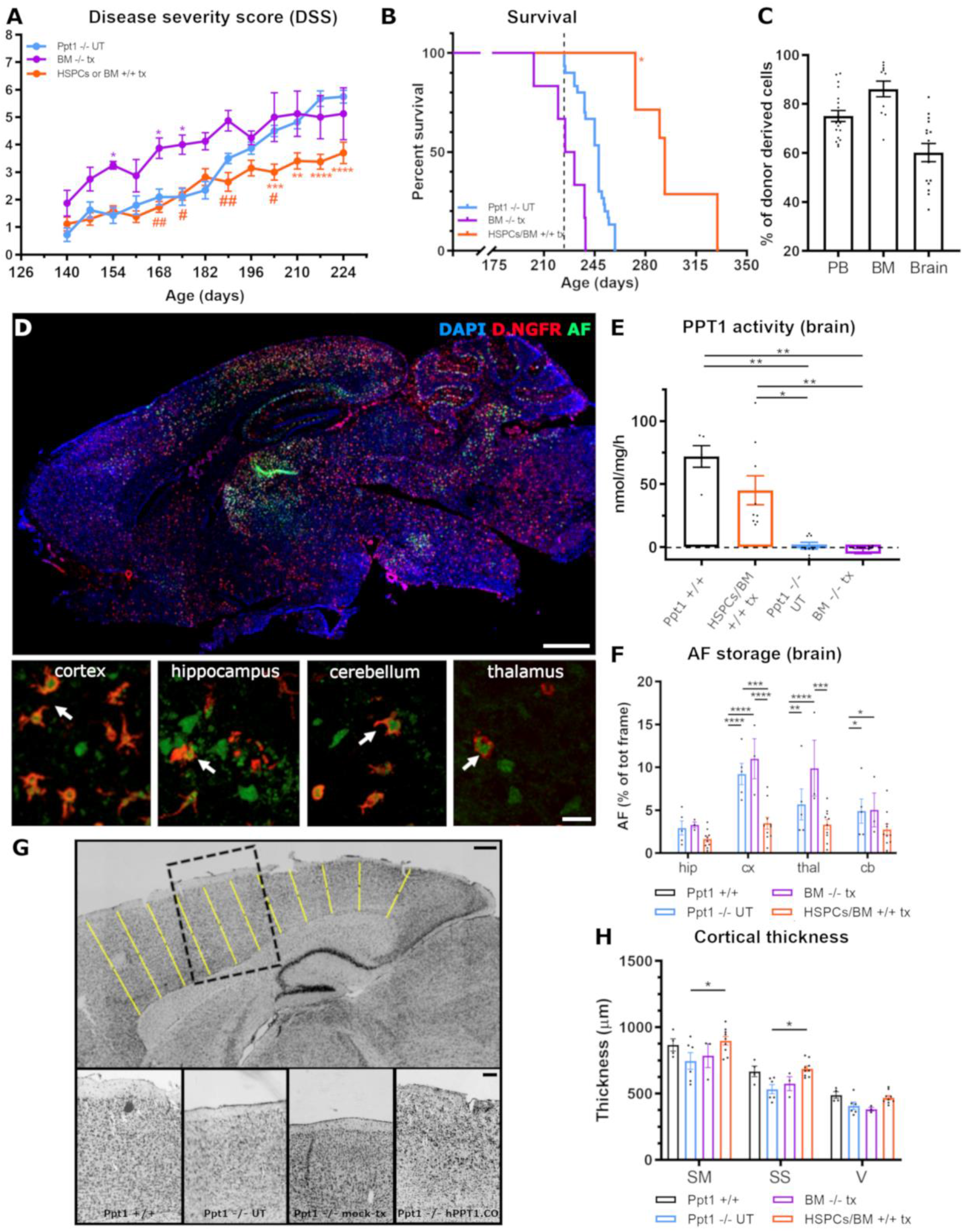
Mild and long-lasting therapeutic effect of wild-type HSPCs/BM transplantation in Ppt1^-/-^ mice. **A**. Ppt1^-/-^ mice transplanted with HSCPs or BM isolated from Ppt1^+/+^ mice display a milder disease progression, assessed by DSS, after onset of the symptoms, as compared to untreated or mock-transplanted Ppt1^-/-^ mice. **B**. Significant increase of survival in Ppt1^-/-^ animals transplanted with HSCPs or BM isolated from Ppt1^+/+^ mice; Log-rank: HSPCs/BM^+/+^ tx vs BM^-/-^ tx p<0.001; BM^-/-^ tx *vs* Ppt1^-/-^ UT p < 0.001. **C**. Histograms showing the donor-cell chimerism in the peripheral blood (PB) at 60 days post-transplantation and in the BM or in the brain upon euthanasia at the end of the study. **D**. Representative fluorescence microscope photomicrographs showing the distribution of donor-derived cells (transduced with D.NGFR reporter gene, red) and autofluorescent storage material (AF, green) in the brain of a Ppt1^-/-^ mouse euthanized at 240 days of age, after transplantation with HSPCs isolated from Ppt1^+/+^ mice. Insets in D show high magnification representative confocal scanning photomicrographs of DNGFR+AF+ cells in different regions of the CNS. **E**. Ppt1 enzymatic activity in the brain of untreated (Ppt1^-/-^) or mock transplanted (BM^-/-^ tx) mice or Ppt1^-/-^ animals transplanted with Ppt1^+/+^ derived HSPCs or BM. The enzymatic activity in wild type Ppt1^+/+^ mice is also reported, as reference. * = p< 0.05; ** = p < 0.01; Kruskal Wallis followed by Dunn’s *post-hoc* test. **F**. Quantification of autofluorescent storage material accumulation in different brain regions (hip=hippocampus, cx=cortex, thal=thalamus and cb=cerebellum). * = p< 0.05; ** = p < 0.01; *** = p< 0.001; ****= p< 0.0001; 2-way ANOVA followed by Tukey’s *post-hoc* test. The histograms of Ppt1^+/+^ group are not visible in the graph as the AF signal was zero in these animals. **G**. Representative brightfield photomicrographs of Nissl-stained brain sections from wild type Ppt1^+/+^ mice and untreated or transplanted Ppt1^-/-^ mice. Black dashed box highlights a portion of the cortex, shown in the high magnification insets. **H**. Histograms showing the quantification of cortical thickness. * = p< 0.05; ** = p < 0.01; *** = p< 0.001; ****= p< 0.0001; Kruskal Wallis followed by Dunn’s *post-hoc* test.

Average donor-cell chimerism, shortly after transplant (in peripheral blood, PB) and at sacrifice (BM and brain), was 80% in PB and BM, and 55% within the brain CD11b^+^ myeloid compartment (Fig. 1C, Fig. S2C,D). Interestingly, there was widespread distribution of cells expressing D.NGFR in the CNS of Ppt1^-/-^ recipients (Fig. 1D), also in areas particularly affected by the pathology, such as the cortex, hippocampus, cerebellum and thalamus (high magnification insets in Fig. 1D). In line with these data, the analysis of PPT1 activity on brain homogenates from mice treated with congenic (Ppt1 +/+) BM or HSPC cells, confirmed a partial reconstitution of the enzymatic activity in the CNS of transplant recipients (Fig. 1E, Fig. S2E).

Autofluorescent (AF) storage accumulation in the CNS is a hallmark of CLN1 pathology, observed in humans and recapitulated in Ppt1^-/-^ mice^5^. Thus, we evaluated and measured the AF material on slices obtained from the brain (cortex, thalamus, hippocampus and cerebellum) of transplanted and control mice sacrificed at the HEP or at 230-237 days of age. Ppt1^-/-^ UT and mock-transplanted mice displayed a striking accumulation of AF material (Fig. 1F), which was absent in WT mice. Treated animals showed an overall reduction of AF (Fig. 1F, S2F). Interestingly, the AF material was also present within the cell body of donor derived cells in the four brain areas (cortex, hippocampus, cerebellum and thalamus) that we analyzed (arrows in Fig. 1D). Based on these results, we measured the cortical thickness on brain sections from treated and control animals colored with cresyl violet (Nissl) staining, as readout of neuronal survival, confirming a rescue of neurons in the somatosensory, somatomotor and visual cortex of mice transplanted with Lin^-^ or total BM WT cells (Fig. 1G,H and Fig. S2G).

### Transplantation of hPPT1-LV transduced HSPCs significantly ameliorates the survival and the phenotype of Ppt1^-/-^ mice

Increasing up to above-normal levels the expression of the WT hydrolases in HSPCs (and their tissue progeny) by LV-transduction represents an efficacious and clinically applicable approach for enhancing therapeutic efficacy of HSPC transplantation in mouse models and patients affected by other LSDs ^24^. Therefore, we tested whether this concept would successfully apply also to CLN1 disease. A 3^rd^ generation LV was generated encoding a codon optimized human PPT1 cDNA under the control of the human phosphoglycerate-kinase (PGK) promoter (hPPT1-LV, Fig. 2A) for transducing Ppt1^-/-^ Lin^-^ HSPCs^20^. The transduced cells (1×10^6 cells/mouse) were transplanted IV in young adult, 40-50 days old, busulfan myeloablated Ppt1^-/-^ recipients. Given the toxicity observed upon usage of Busilvex™ in previous transplants, for these experiments we adopted a lab-grade (more tolerable) formulation of busulfan dissolved in acetone and peanut-oil^25^. As control groups, myeloablated Ppt1^-/-^ mice transplanted with cultured un-transduced Ppt1^-/-^ HSPCs and untreated Ppt1^-/-^ animals were employed. Myeloablated Ppt1^+/+^ WT animals transplanted with Ppt1^+/+^ HSPCs or untreated Ppt1^+/+^ mice were also included as reference during behavioral and biochemical assessments. After transplant, animals were monitored for survival and disease-associated phenotype until they reached the HEP or up to study termination.

**Fig 2.**
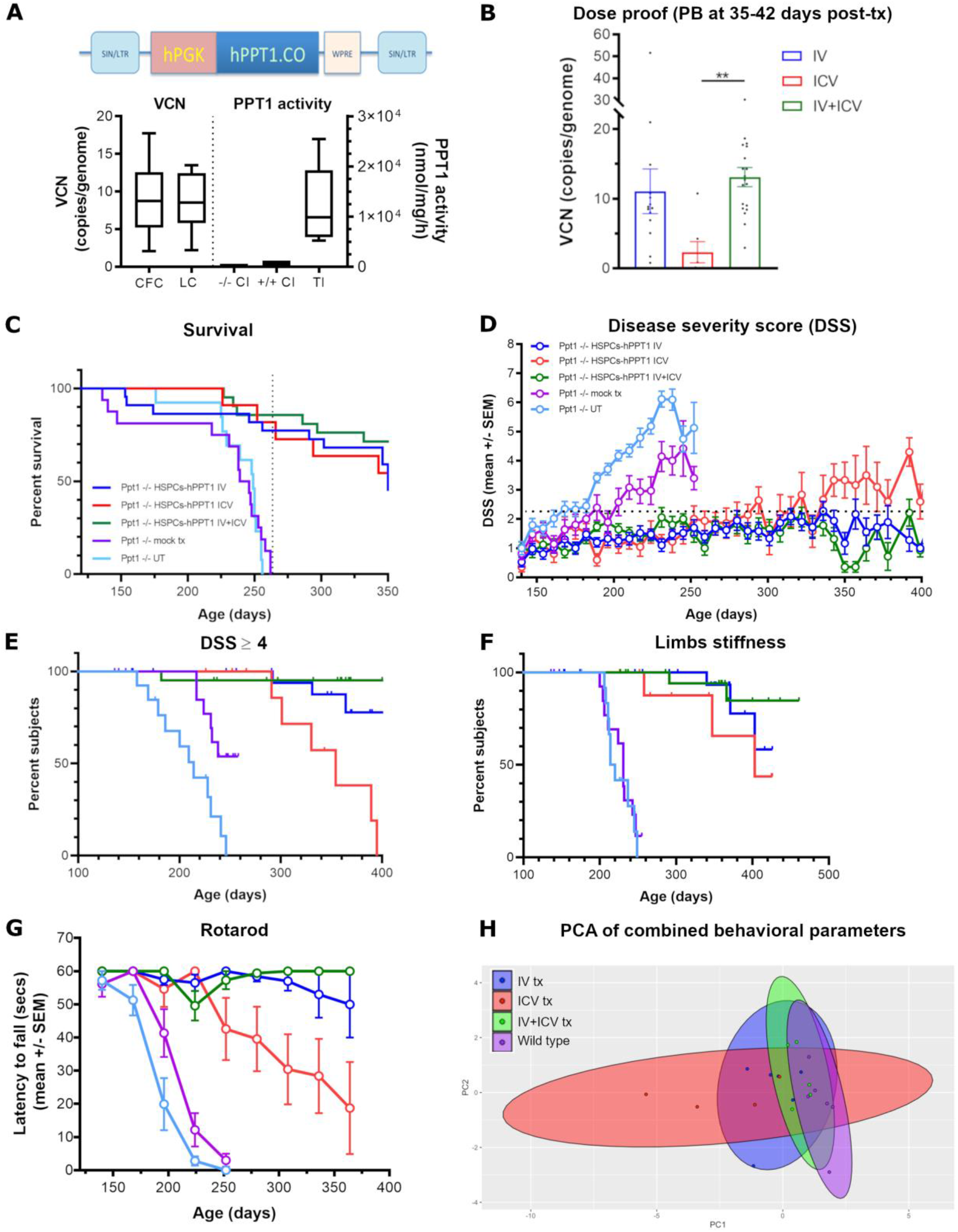
Prevention of neurological deficits and increased survival of Ppt1^-/-^ mice transplanted with hPPT1-LV transduced HSPCs. **A**. High transduction efficiency of HSCPs assessed in colony forming units and suspension cultures at 14 days after transduction with a lentiviral vector expressing codon optimized human PPT1 (left graph). Supraphysiological Ppt1 enzymatic activity in hPPT1-LV transduced HSPCs as compared to Ppt1^+/+^ or Ppt1^-/-^ mock transduced HSPCs (right graph). **B**. High donor cell chimerism in peripheral blood (PB) of mice transplanted IV or IV+ICV with hPPT1-LV transduced HSPCs. ** = p < 0.01. Kruskal Wallis followed by Dunn’s post-hoc test. **C**. Significant increase of the survival in Ppt1^-/-^ mice receiving the administration of hPPT1-LV transduced HSPCs. **D**. Long-lasting prevention of disease manifestations in Ppt1 ^-/-^ mice transplanted with hPPT1-LV transduced HSPCs up to the study termination (around 360-400 days). **E, F**. Significant prevention of pathological symptoms (DSS higher than 4 and limbs stiffness) in Ppt1^-/-^ mice transplanted with hPPT1-LV transduced HSPCs. **G**. Significant prevention of the deterioration of motor performances (assessed with rotarod) in Ppt1^-/-^ mice transplanted with hPPT1-LV transduced HSPCs. **H**. Principal component analysis of combined behavioral data. See supplementary table 3 for statistics.

A high transduction efficiency of the Ppt1^-/-^ HSPCs was obtained with the newly developed LV (9 vector copies/genome on average), as measured on the output of the colony forming assay (CFC) or of the liquid culture (LC) (assessed at 14 days after transduction, Fig. 2A, left panel) of the batches used for n=15 transplantations. This corresponded to an above-normal PPT1 activity in the LC from hPPT1-LV transduced Ppt1^-/-^ HSPCs, that reached ∼20-fold the enzymatic activity measured in the LC derived from Ppt1^+/+^ HSPCs (Fig. 2A, right panel). Dose proof was obtained 35-42 days after transplant, by demonstrating high vector copy number (VCN) in the PB of transplanted animals (Fig. 2B).

Animals transplanted intravenously (IV) with the hPPT1-LV transduced HSPCs displayed a significantly increased survival as compared to UT or mock-transplanted Ppt1^-/-^ control mice. Indeed, while all the UT and mock-transplanted animals reached HEP or were in poor body conditions (PBCs) due to the pathology by 260 days of age (Fig. 2C), 77% of the gene therapy treated animals (17 out of 22) were still alive at 260 days, and 68% of the initial cohort (15 out of 22) survived until the study was terminated at ≥350 days of age (Fig. 2C). The recorded intercurrent deaths (ICDs) were mainly attributable to animals displaying PBCs in the absence of clinically relevant symptoms associated with CLN1 disease. This was further investigated by necropsy and histopathological assessments and the ICDs were generally not attributed to disease progression (Fig. S4B), but rather to conditioning related effects^26^.

Behavioral parameters were recorded on treated and control mice from 140 days of age until sacrifice (with a window of study termination that lasted from 350 days to 500 days of age). Importantly, the weekly DSS assessment and monthly rotarod testing demonstrated that the majority of the gene therapy treated animals were protected from the development of severe neurological symptoms. Indeed, while UT and mock-transplanted animals experienced a progressive neurological deterioration after the age of 220 days, with half of them in disease stage 2 (DSS ≥ 4) at 220-240 days of age (Fig. 2D-E), the majority of the transplanted mice showed no clinical onset of the disease (DSS ≤ 2) for the whole duration of the observation (Fig. 2G). Of the entire cohort of 22 treated mice, only 4 mice (18%) showed a mild pathology (DSS ≥ 4, Fig. 2D-E) at study termination (by 350-360 days), and only 3 animals experienced a worsening of symptoms, identified as hindlimbs stiffness and 50% reduction of rotarod performance, after 320 days of age (Fig. 2F,G).

### Intra-CNS HSC gene therapy attenuates disease symptoms in Ppt1^*-/-*^ mice

We recently demonstrated that transplanting HSPCs directly in the cerebral lateral ventricles of busulfan-myeloablated recipients fosters reconstitution of the brain myeloid compartment by the transplanted cell progeny, leading to rapid, CNS-restricted engraftment of transplant-derived microglia-like cells^27^. Thus, we assessed the therapeutic potential of hPPT1-LV transduced HSPC administration directly in the cerebral lateral ventricles of myeloablated Ppt1^-/-^ recipients as a single treatment. This was done to understand whether a CNS-restricted engraftment of the transduced, PPT1 expressing cells could be sufficient to alleviate CLN1 disease manifestations. Ppt1^-/-^ Lin^-^ HSPCs, transduced as described above with the PPT1 encoding LV, were transplanted (0.3×10^6 cells/mouse) in the CNS by monolateral intracerebroventricular (ICV) injection in young adult, 40-50 days old, busulfan myeloablated Ppt1^-/-^ mice. Un-manipulated Ppt1^-/-^ BM cells were administered 5 days after ICV cell transplantation to aid hematopoietic rescue, as we do not expect significant contribution to hematopoiesis (no BM engraftment) of the ICV-injected cells^27^.

Animals treated by this CNS-directed approach (tot n=10) showed an early benefit from the treatment, as 80% of them were alive and devoid of neurological symptoms (DSS<2) at around 260 days of age, when all UT and mock treated mice had already reached HEP (Fig. 2C). However, later-on, their behavioral assessment highlighted onset of symptoms and they became clearly symptomatic with a DSS ≥ 4 (Fig. 2D, E) and progressive motor deficits at rotarod (Fig. 2G) by 300 days of age. In 3 out of 10 animals from this group, hindlimbs stiffness was also observed starting from 340 days of age (Fig. 2F) and a fraction of them reached HEP due to the pathology before study termination. By approximately 325 days of age their survival dropped to 60% (Fig. 2C).

### Combined IV and ICV delivery of the transduced HSPCs enhances the efficacy of gene therapy leading to complete prevention of CLN1 disease symptoms

The severe and rapid progression of CLN1 disease in patients imposes the need for rapid and extensive restoration of the enzymatic activity in the CNS to increase the likelihood of preventing and/or halting disease manifestations. Based on the findings reported above, we tested the role of transduced HSPC delivery in the CNS as an additive strategy to enhance HSC gene therapy efficacy in a combinatorial (IV+ICV) transplant setting. This combined approach was intended at fostering and enhancing the myeloid CNS engraftment of the transplanted cells, the delivery of therapeutic enzyme to the CNS of Ppt1^-/-^ recipients and the overall therapeutic potential of our treatment. Ppt1^-/-^ Lin^-^ HSPCs transduced with hPPT1-LV were transplanted in 40-50 days old, busulfan myeloablated Ppt1^-/-^ recipients IV and ICV, as detailed above, in the same day, ≥ 24 hours after the last busulfan dose. As control group, myeloablated Ppt1^-/-^ mice IV-and ICV-transplanted with cultured un-transduced Ppt1^-/-^ HSPCs were employed.

Interestingly, the combinatorial ICV+IV treatment resulted in a robust therapeutic benefit with complete prevention of disease manifestations in the large majority of the treated mice. Indeed, 86% of the ICV+IV transplanted animals (tot n=21) were still alive and devoid of any neurological symptoms (Fig. 2C,D) at around 260 days of age. 76% of them outlived controls and were sacrificed for study termination at 400-460 days of age with no evidence of disease manifestations (DSS ≤ 2, no limb clasping nor stiffness, preserved muscle tone) and full performance at rotarod test (Fig. 2E,F,G).

Importantly, by applying principal component analysis (PCA) on the multiple behavioral parameters measured in each group (namely, survival, DSS and rotarod performance at 230, 330 and 350 days of age) we highlighted a close similarity between animals transplanted with hPPT1-LV HSPCs IV+ICV and WT mice. In contrast, results for animals transplanted with hPPT1-LV HSPCs via the IV-only or ICV-only routes showed a more scattered distribution on the PCA plot. Multivariate analysis of variance and Hotelling’s T-test confirmed a significant difference between animals transplanted IV+ICV and the animals that received the cells only IV (p<0.05) or ICV (p<0.01), suggesting that the combinatorial transplant strategy could achieve greater (and more homogeneous across subjects) clinical benefit than the individual approaches.

### HSC gene therapy provides biochemical and pathological rescue of CLN1 disease: group comparison reveals an advantage of ICV+IV gene therapy over the other treatment approaches

To support the behavioral findings reported above, we investigated the CNS tissues of the gene therapy treated and control animals from each group, collected at study termination or when the mouse was euthanized. A robust engraftment of the transplanted cells was documented in the CNS of the gene therapy recipients, as measured by quantification of integrated hPPT1-LV by ddPCR (vector copy number per cell) in the brain and spinal cord (Fig. 3A). A higher VCN was measured in IV and IV+ICV groups as compared to ICV only, in both brain and spinal cord. Importantly, a remarkable reconstitution of PPT1 activity was measured in the brain and spinal cord of gene therapy recipients, with 2-to 4-fold increase of enzyme activity above the physiological levels of Ppt1^+/+^ mice (Fig. 3B). The highest PPT1 activity levels were measured in the brain of IV+ICV-transplanted mice (Fig. 3B). Lower PPT1 activity was measured in the spinal cord, but not in the brain, of ICV versus IV and IV+ICV-transplanted mice (Fig. 3B). IV and IV+ICV transplanted mice also had successful transduced cell engrafment (as measured by VCN) and robust PPT1 enzyme activity reconstitution in the BM (Fig. S4A). As expected, negligible raise in PPT1 activity and transduced cell engraftment were measured in the BM of ICV-transplanted animals, with the exception of three animals displaying detectable VCN and PPT1 activity (Fig. S4A) suggesting partial leakage from the ICV transplantation These findings correlated with evidence of sustained metabolic correction in the CNS of the treated animals. Indeed, treated animals showed a clear-cut prevention of AF storage accumulation in the cortex, thalamus and hippocampus, whereas in the cerebellum storage material was not reduced for animals transplanted ICV (Fig. 3C,D). In particular, in Ppt1^-/-^ UT and ICV transplanted mice we observed accumulation of fluorescent material in the few surviving Purkinje cells (arrowheads in Fig. 3C). In contrast, IV and IV+ICV transplanted mice displayed less prominent fluorescent material in Purkinje cells (arrows in Fig. 3C). Consistent with these results, Ppt1^-/-^ mice transplanted with hPPT1-LV HSPCs via IV or IV+ICV routes displayed a significant protection from neuronal demise, resulting in a cortical thickness comparable to WT mice and significantly different from UT and mock-transplanted Ppt1^-/-^ mice (Fig. 4A). Notably, ICV-only transplanted animals showed a tendency to amelioration without reaching statistical significance (Fig. 4A).

**Fig 3.**
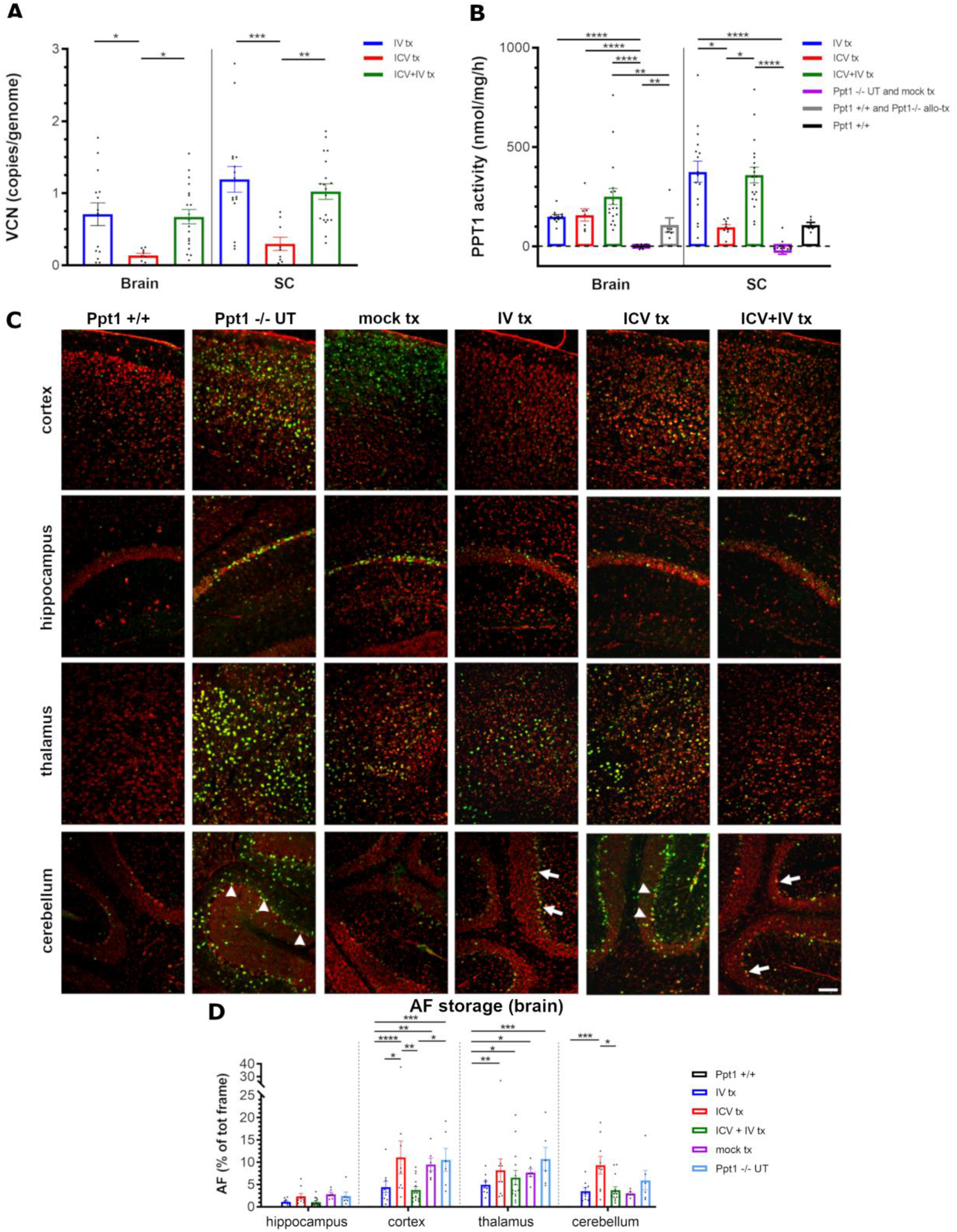
Rescue of PPT1 activity and prevention of autofluorescence storage accumulation in the brain of Ppt1^-/-^ mice transplanted with hPPT1-LV transduced HSPCs. **A, B**. Donor-cell chimerism (VCN) and Ppt1 enzymatic activity assessed in brain and spinal cord of untreated or transplanted mice analyzed at study termination. * = p < 0.05; ** = p < 0.01; *** = p < 0.001; **** = p < 0.00001; Kruskal Wallis followed by Dunn’s post-hoc test. **C**. Representative fluorescence microscope photomicrographs showing autofluorescent storage material (AF, green) in different brain regions of Ppt1^-/-^ mice transplanted with hPPT1-LV transduced HSPCs at 400-420 days of age, when the study was terminated. Mock transplanted and untreated Ppt1 ^-/-^ mice at about 250 days, i.e. humane end point, are shown as reference; 450 days-old untreated wild type Ppt1^+/+^ are shown as control. Fluorescent Nissl staining (red) is used to highlight neurons in different brain regions. **D**. Quantification of autofluorescent (AF) storage material in different brain regions of untreated or transplanted mice analyzed at study termination. * = p < 0.05; ** = p < 0.01; *** = p < 0.001; **** = p < 0.00001; 2-way ANOVA followed by Tukey’s post-hoc test.

**Fig 4.**
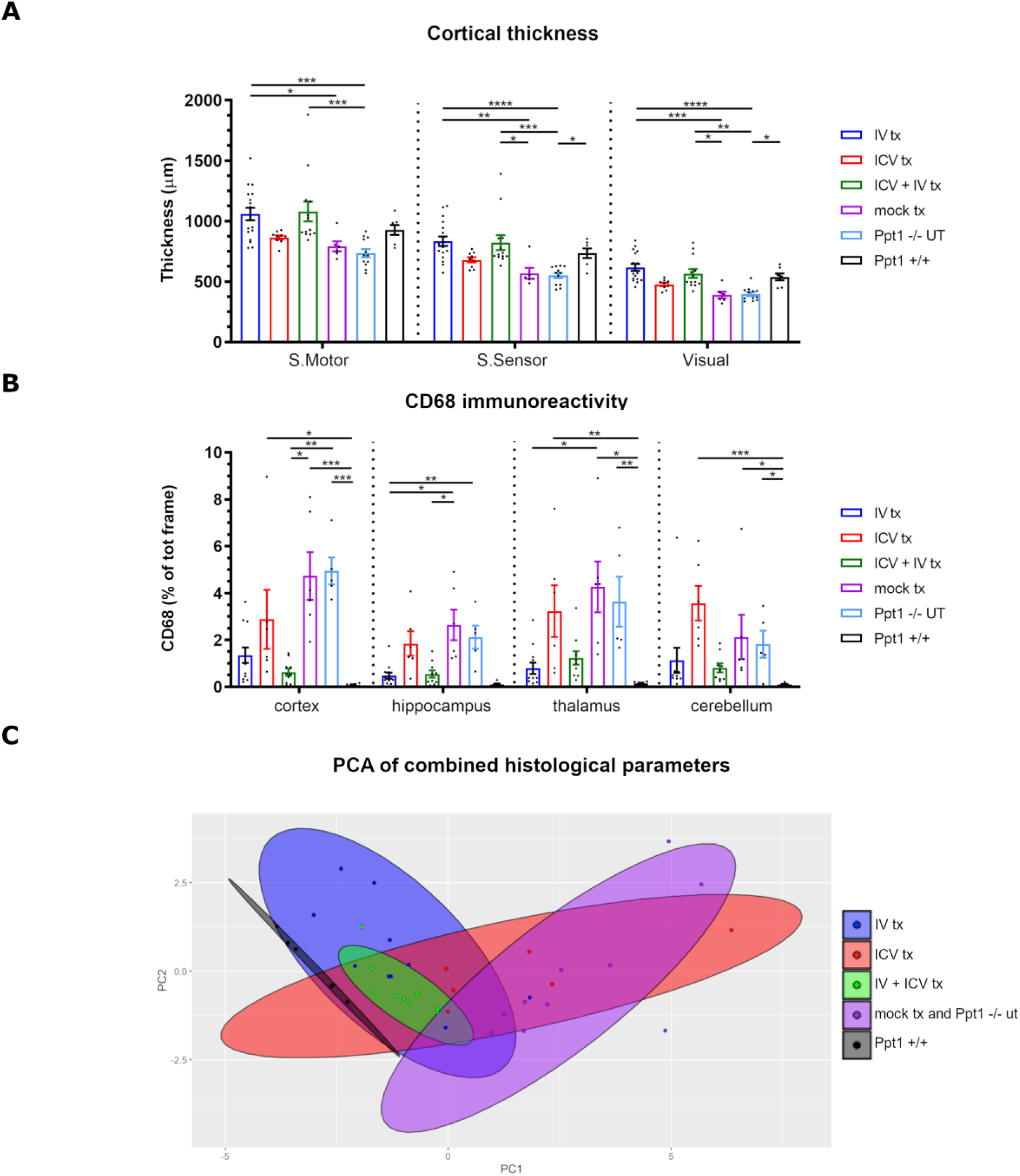
HSC gene therapy rescues neuronal survival and mitigates microgliosis in Ppt1 ^-/-^ mice. **A**. Quantification of cortical thickness in the somatosensory (SS), somatomotor (SM) and visual (V) area, in untreated Ppt1^-/-^ or Ppt1^+/+^ mice and in mice transplanted (tx) with hPPT1-LV transduced HSPCs IV, ICV or ICV+IV or mock-transplanted, analyzed at study termination. * = p < 0.05; ** = p < 0.01; *** = p < 0.001; **** = p < 0.00001; 2-way ANOVA followed by Tukey’s *post-hoc* test. **B**. Quantification of CD68 immunoreactivity in the cortex, hippocampus, thalamus and cerebellum of untreated (UT) or gene therapy treated Ppt1^-/-^ mice. * = p < 0.05; ** = p < 0.01; *** = p < 0.001; **** = p < 0.00001; 2-way ANOVA followed by Tukey’s *post-hoc* test. **C**. Principal component analysis of combined histopathological data.

Neuroinflammation is a hallmark of pathology in CLN1 disease^12,16,17^. Early and progressive activation of microglia and astrocytes have been described in the Ppt1^-/-^ mice, in brain areas critically affected by storage accumulation and neuronal demise. We investigated how the engrafted donor-derived cells affected the extent of glial reactivity by quantifying CD68 as a marker of microglia/macrophages. As expected, extensive CD68 immunoreactivity (characterized by many CD68^+^ ameboid-like cells scattered throughout the parenchyma) was detected in the cortex, hippocampus, thalamus and cerebellum of end stage Ppt1^-/-^ and mock-transplanted mice (arrows in Fig. S3 and quantification in Fig. 4B), consistent with the massive accumulation of AF storage material and severe neuronal loss that we reported in these regions (Fig. 3C,D). In contrast, generally very few CD68^+^ cells were detected in the same regions of hPPT1-LV HSPCs treated mice. Again, animals receiving the cells via either IV or IV+ICV routes displayed the greatest effect, with only few small CD68^+^ cells distributed sparsely throughout the parenchyma (Fig. S3). IV+ICV animals showed the lowest CD68 immunoreactivity of the entire analyzed cohort in the cortex (Fig. 4B). A more pronounced CD68 immunoreactivity was instead observed in animals injected with hPPT1-LV HSPCs by the ICV-only route (Fig. 4B), especially in the thalamus and cerebellum (arrowheads in Fig. S3), suggestive of an ongoing reactive gliosis in line with the AF storage accumulation and neuronal loss observed in these animals.

PCA was also applied on the multiple described histological parameters measured in each group (namely, AF storage and CD68 immunoreactivity in cortex, thalamus, hippocampus and cerebellum, and thickness of the cortical layer) with the aim of assessing the extent to which the different hPPT1-LV HSPC delivery routes affected these parameters. As shown in Fig. 4C, animals transplanted with IV and IV+ICV hPPT1-LV HSPCs overall clustered in the proximity of Ppt1^+/+^ healthy mice and far from mock-transplanted and untreated Ppt1^-/-^ mice. However, while ICV+IV transplanted mice showed the most homogenous clustering, suggestive of low variability, IV transplanted mice showed scattered individuals. Multivariate analysis of variance and Hotelling’s T-test confirmed that gene therapy treated Ppt1^-/-^ mice are significantly different from the Ppt1^-/-^ UT group; interestingly, among the gene-therapy groups, animals transplanted IV+ICV are significantly different from the IV (p < 0.05) and ICV (p < 0.001) groups. This further supports the evidence that a stronger and more robust rescue of the pathological CLN1 disease phenotype is obtained when hPPT1-LV HSPCs are transplanted IV+ICV, a treatment protocol that provided a homogeneous beneficial effect among the different treated individuals. In the case of ICV gene therapy, the efficacy was only partial, especially in the long-term observation.

### Combined intra-CNS and IV gene therapy prolongs survival and alleviates the phenotype of symptomatic Ppt1^*-/-*^ mice

Based on the information gained from this comparative study, we decided to treat with the hPPT1-LV HSPCs (administered IV or through the combined IV+ICV approach) Ppt1^-/-^ mice at onset of symptoms, at 130±15 days of age (Fig. S1B), to verify whether we could achieve therapeutic benefit also in a stage of the disease when damage has already accumulated in the CNS. Animals received the standard pre-transplant busulfan conditioning regimen and anti-seizure prophylaxis and the same dose of hPPT1-LV transduced HSPCs as the young-adult mice that were transplanted in the previous experiments. To generate additional feasibility and toxicology data, in parallel to the gene therapy-treated Ppt1^-/-^ animals, we also generated two large groups of mock-transplanted, Ppt1^-/-^ and Ppt1^+/+^ animals (n=21 and n=24, respectively), which were exposed to the same busulfan conditioning regimen, but transplanted with untransduced HSPCs (from Ppt1^-/-^ or Ppt1^+/+^ donors, respectively). Symptomatic mice transplanted IV or IV+ICV with the transduced HSPCs survived significantly longer than Ppt1^-/-^ UT and mock-transplanted mice, with about 80% of the treated animals still alive at 260 days of age, when all animals from Ppt1^-/-^ control groups had already died. Survival subsequently dropped also in treated mice, but the IV+ICV group showed a statistically greater survival probability than the IV one up to study termination (Fig. 5A), indicating a more robust effect of gene therapy when the transduced cells were delivered also in the CNS. This was consistent with a significant prevention of late disease manifestations, as confirmed by the behavioral assessment. Despite the worsening of symptoms, as measured by the DSS, of gene therapy treated animals apparently progressed broadly parallel to Ppt1^-/-^ UT and mock transplanted mice at least up until 250 days of age (Fig. 5C), after this stage the pathology stabilized around a DSS score of ∼ 4.5. Notably, 80% of treated animals never displayed hindlimbs stiffness (Fig. 5C) which represent the most evident symptom displayed by Ppt1 -/- animals when they reach the advanced stage of the pathology. Consistent with these data, the analysis of donor cell engraftment (measured as vector genome copies) performed at study termination confirmed high VCNs both in peripheral organs and in the CNS of the treated mice (Fig. 5B, left graph). Interestingly, VCN in the BM and, most importantly, in the brain and spinal cord of IV+ICV treated mice was higher than the VCN measured in IV transplanted animals (Fig. 5B). Accordingly, very high, above-normal PPT1 activity levels were detected in the CNS and BM of gene therapy treated mice (Fig. 5B). Of note, also in this case IV+ICV cell transplantation was associated to an higher enzymatic activity in the brain (and BM) as compared to IV treated animals (Fig. 3B, 5B), confirming the advantage of combined cell delivery.

**Fig 5.**
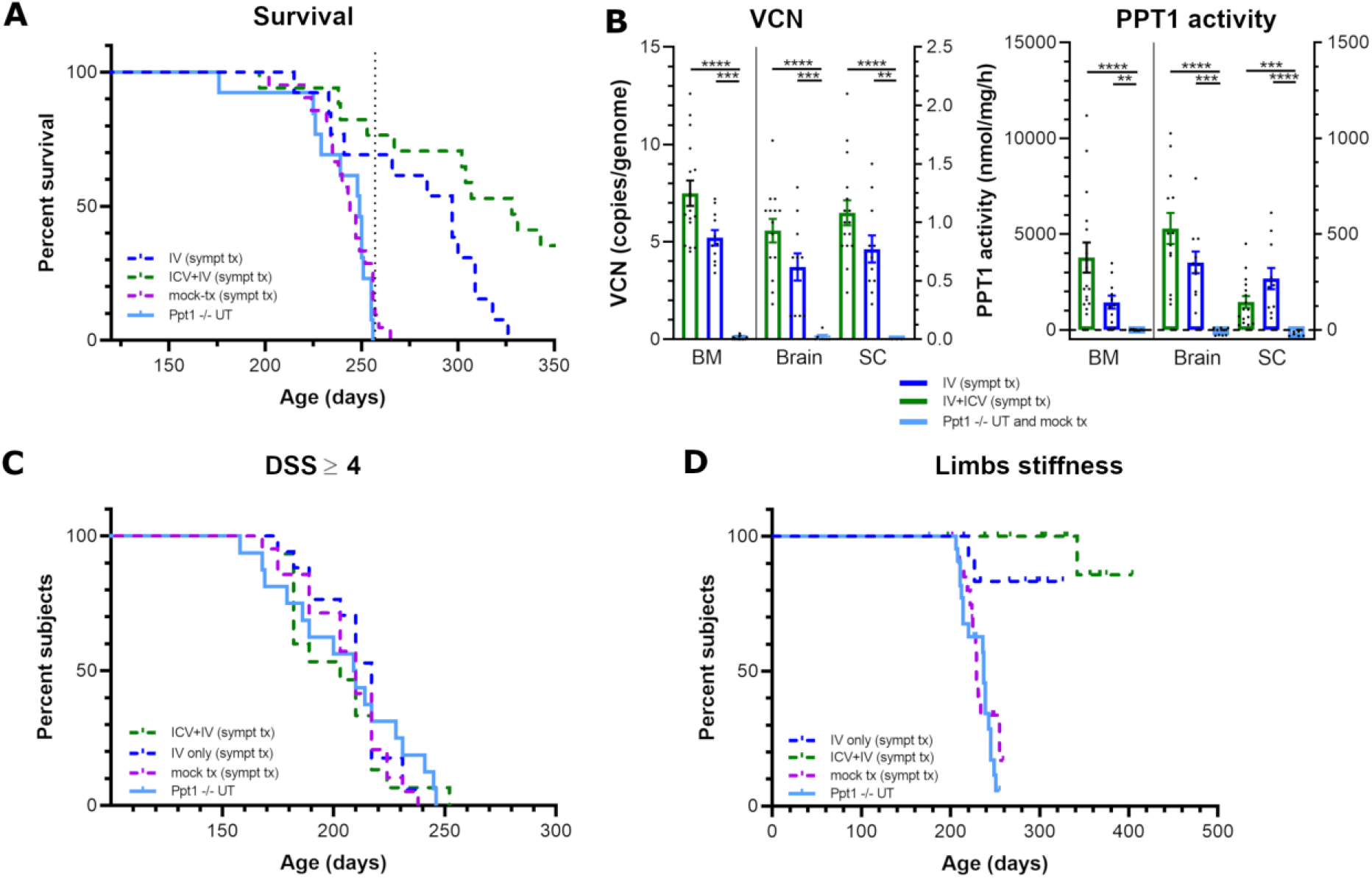
Clinical benefit of HSC gene therapy in Ppt1^-/-^ mice transplanted at the symptomatic stage. A. Significant increase of the survival in Ppt1^-/-^ mice transplanted at the symptomatic stage with hPPT1-LV transduced HSPCs administered IV+ICV. B. High donor cell chimerism (vector copy number – VCN -, left panel) and increase of PPT1 enzymatic activity (right panel) in bone marrow (BM), brain and spinal cord of mice transplanted with hPPT1-LV transduced HSPCs. *** = p< 0.001; **** = p< 0.0001; Kruskal Wallis followed by Dunn’s post-hoc test. **C**. Similar progression of symptomathology in all treated groups until about 250 days of age. **D**. Significant delay of the deterioration of motor performances (hindlimb stiffness) in Ppt1^-/-^ mice transplanted with hPPT1-LV transduced HSPCs. See supplementary table 3 for statistics.

### Safety of hPPT1-LV HSPCs transplantation in Ppt1^-/-^ mice

To collect information on the safety of transplantation of hPPT1-LV HSPCs in Ppt1^-/-^ mice, the follow-up monitoring of the different experimental groups (treated, UT and mock-transplanted PPT1^-/-^ and WT animals) included a large set of assessments. Among them, we monitored post-transplant and long-term hematopoiesis by evaluation of PB composition at 5-6 weeks post-transplant and at time of euthanasia, to verify the repopulation and multilineage differentiation potential the transplanted HSPCs and to exclude the presence of clonal alterations or other hematopoietic anomalies. Overall, at the first time-point no significant differences were observed between all the groups, suggesting that hematopoietic reconstitution occurred normally for all treated groups irrespective of the type of cells transplanted and the route of administration (Fig. S4C). At euthanasia, normal PB composition was also documented, with absence of hematopoietic anomalies in the treated and control groups (Fig. S4D).

The clinical follow up of treated and control animals consisted of twice weekly monitoring for viability and health status. Any intercurrent death (animal found dead or requiring euthanasia due to poor body condition) was registered and full necropsy was performed. As shown in Fig. S4B, while the majority of Ppt1^-/-^ animals from the mock-transplanted and UT groups underwent euthanasia because they reached the pathological HEP, none of the Ppt1^-/-^ mice that were transplanted with hPPT1-LV HSPCs IV or ICV+IV had to be euthanized due to CLN1 disease symptoms. About 20% of animals from ICV-only group did require euthanasia for reaching the HEP. We reported in total 34 animals (spread across all groups, including mock-transplanted PPT1^-/-^ and WT mice, except the Ppt1^-/-^ UT) that had to be euthanized due to poor body conditions likely caused by abdominal ascites (confirmed by necropsy observation) and frequently accompanied by the presence of a solid tumor mass in the abdominal cavity (Supplementary Table 4). Histopathological examination identified the tumor masses as sarcomas accompanied by atypical fibroplasia of the abdominal cavity. These sarcomas were attached to, and appeared to arise from, serosal/peritoneal surfaces (Supplementary Table 5). One mass, that was observed in Ppt1^+/+^ un-transplanted animals, correlated with lymphoma and was considered to be related to a spontaneous tumor event that is reported to occasionally occur in wild type C57BL/6J animals after 300 days of age.^28,29^ The sarcomas were all similar in appearance and were diagnosed as sarcoma NOS (not otherwise specified) because they lacked collagen and were generally fairly pleomorphic. These findings were observed only in the groups that underwent busulfan conditioning, that is administered intraperitoneally, followed by transplantation of HSPCs (either transduced or untransduced), and they occurred more frequently in animals that managed to survive longer than 260 days. In line with our observation, the toxicity associated with the route of administration of busulfan that in mice must be intraperitoneal for technical reasons, was reported in the context of other long-term observation preclinical studies ^26^.

The histopathological analyses of hematopoietic organs, including spleen and BM, showed that Ppt1-/- UT mice had decreased spleen size (subgross), decreased white pulp cellularity and increased apoptosis, macrophage aggregates and adipocytes when compared to Ppt1+/+ mice; whereas in the BM, they had increased macrophage cellularity, suggesting that these findings represent overall a feature of the Ppt1-/- strain. These findings were similarly observed in mock-transplanted Ppt1-/- and in ICV treated mice, whereas they were reduced in the animals that received hPPT1-LV transduced HSPCs IV or ICV+IV, supporting the beneficial effect exerted in the peripheral tissues by the functional hydrolase ^15^ released by the systemically transplanted hPPT1-expressing HSPCs. No hematopoietic clonal proliferations were observed.

## Discussion

CLN1 disease is one of the most aggressive forms of NCL still lacking a curative treatment. We and others demonstrated that HSC progeny cells can represent a vehicle for therapeutic molecule delivery to the CNS and exert neuroimmunomodulatory functions in the CNS upon transplantation in myeloablated host^20,24,30^. For this reason, we here investigated whether HSC transplantation (HCT) and/or HSC gene therapy could determine widespread restoration of Ppt1 enzymatic activity throughout the nervous system and provide therapeutic benefit in the CLN1 disease animal model. To favor the establishment of a high donor-cell chimerism in the recipients’ CNS^22,31^, we employed a busulfan-based conditioning regimen that is instrumental to foster efficient engraftment of donor-derived cells post-transplant^25^. To accurately monitor the effects of our proposed therapeutic treatment, we developed a new clinical scoring system (called disease severity score -DSS) that enables precise highlighting of early signs of the disease and monitoring of the progressive behavioral deterioration of untreated Ppt1^-/-^ mice in parallel with other tests, such as the open field test^10^ or the rotarod (our work and ref.^22^), by which deficits become significantly evident only at advanced disease stages when animals are already severely compromised.

Interestingly, standard WT HSC transplantation resulted in an unexpected clinical benefit. Indeed, transplant recipients exceeded in survival the natural disease course of Ppt1^-/-^ UT mice and developed only a mild disease phenotype, up until the age of death of the first mock-transplanted control (224 days), with a mild DSS trajectory over time. *Ex vivo* analyses demonstrated a high donor-cell chimerism both in hematopoietic cells and in brain CD11b^+^ myeloid/microglia cell subset. At the histological level, we observed widespread distribution of the progeny of the transplanted cells throughout the brain parenchyma, including in the regions particularly susceptible to the disease, where the donor cells showed an activated phenotype and appeared engulfed with AF storage material. A partial restoration of Ppt1 enzymatic activity was observed in the brain of these animals at about 50% of the levels found in WT mice. This was sufficient to reduce (but not abrogate) the burden of AF material accumulation (especially in the thalamus and cortex) and prevent the thinning of the cortical layer, a hallmark of neurodegeneration in CLN1 disease. Notably, these data represent the first demonstration that HSC transplantation could be beneficial in CLN1 disease, provided that an optimized conditioning regimen is applied. The extension of survival and the rescue of histopathological disease hallmarks observed in the transplant recipients are similar, in extent, to the results described by other groups upon intra-cerebral injection of Ppt1 expressing adeno-associated viral (AAV) vectors^12,22,32^. Notably, we transplanted 40-50 days old Ppt1^-/-^ mice, when early signs of the pathology already started to appear (at least at the histopathological level). This suggests that the HSC-based approach retain a great therapeutic potential also when it is applied at the early stages of the neurodegeneration. Importantly, the evidence of donor-derived cells engulfed with storage material in several brain regions of the transplant recipients also suggests that the cell-therapy approach could determine a benefit through a synergistic effect provided by the engagement of the HSC progeny not only in the release of the functional enzyme but also in the scavenging of toxic debris and storage from the environment.

Based on these findings, we hypothesized that gene transfer could increase the overall therapeutic potential of HCT also in CLN1 disease, as seen in other LSDs ^20,33^. Indeed, multicopy LV gene transfer into Ppt1^-/-^ HSPCs could allow expression of Ppt1 enzyme above physiological level in the transplanted cell progeny in tissues, including the nervous system, and therefore determine more robust clinical benefit. To address this hypothesis, busulfan myeloablated Ppt1^-/-^ young adult recipients were transplanted IV with Ppt1^-/-^ HSPCs efficiently transduced with hPPT1-LV. This approach resulted not only in a significant increase of the survival, but also in a clearcut rescue of the CLN1 disease phenotype in the majority (approx. 80%) of the treated animals up to study termination. In line with this observation, we reported supra-physiological levels of Ppt1 enzymatic activity in the brain and spinal cord of the transplanted mice (∼3.5-fold higher than the WT or WT HSPC transplanted animals). Cortical thinning was prevented, and AF storage accumulation was also reduced in the treated mice, as compared to UT or mock-transplanted controls. The direct comparison of these results with the outcome of WT cell transplant clearly showed the superiority of gene therapy in preventing/arresting clinical and histopathological disease manifestations. Thus, we proved in the preclinical setting that the strategy that led to successful application of HSC gene therapy in children affected by other LSDs with primarily demyelinating features ^24,34^ could be efficacious also in CLN1 disease, that is a purely neurodegenerative condition.

In the attempt of enhancing further the therapeutic potential of HSC gene therapy for neurometabolic conditions and possibly extending treatment indication, we recently developed a novel approach based on the direct administration of the transduced HSPCs in the cerebral lateral ventricles (ICV delivery) of transplant recipients. This route *per se* or in combination with the standard IV cell-delivery accelerates and fosters microglia reconstitution in busulfan conditioned mice^27^. This could represent a suitable strategy to anticipate timing of clinical benefit in neurometabolic diseases, especially for those pathologies like CLN1 disease where CNS damage is rapidly progressive and key to disease course. We thus tested the therapeutic efficacy of the transplantation of hPPT1-LV transduced HSPCs ICV alone or in combination with IV delivery into busulfan myeloablated Ppt1^-/-^ adult recipients. Interestingly, when the transduced HSPCs were delivered exclusively ICV, albeit with un-transduced Ppt1^-/-^ hematopoietic cells transplanted IV to rescue the BM from myeloablation, we observed a significant therapeutic benefit, with increased survival and prevention of disease symptoms up to 250 days of age, when all the animals from the UT or mock-transplanted group had already died. This novel finding supports the concept that ICV transplantation of HSPCs could provide a significant benefit to CNS pathology and associated clinical deficits, a finding of potential value for other disorders where the CNS is the only affected tissue. However, the animals subsequently began to display some symptoms of CLN1 disease, including hindlimbs clasping and stiffness, and rotarod deficits. These deficits progressively worsened, but remained mild (DSS ∼4), in approximately 40% of the treated animals. Moreover, despite histological and biochemical analyses at study termination showed restoration of enzymatic activity at WT levels in the brain and in the spinal cord of transplant recipients, AF storage was not homogenously reduced as compared to control animals, and cortical thinning was not widely prevented. This partial benefit could be interpreted in light of different considerations. Firstly, even if CLN1 disease is considered a LSD with predominant CNS involvement, extra-CNS tissue alterations have been reported in CLN1 disease patients and mice, such as cardiac dysfunction^6^ and metabolic deficits^35^. The accumulating deficits in these tissues are rarely detected due to the overwhelming symptomatology induced in the brain and spinal cord. We demonstrated that ICV transplanted cells mainly repopulate brain and spinal cord with a microglia-like progeny, with little to no engraftment in the extra-CNS and hematopoietic tissues. Thus, we can hypothesize that the ICV approach could have contributed to some rescue of the CNS pathology, as seen by the partial and initial effects, but not to halting extra-CNS manifestations that, once the lifespan was extended by the treatment, became evident and affected the disease course. This was also evidenced by histopathology performed on hematopoietic organs. Moreover, we cannot exclude that partial exhaustion of the ICV transplanted cells and/or of their progeny could have occurred in the long term post-transplant. Indeed, at study termination we detected lower chimerism and Ppt1 activity in the brain and spinal cord of ICV-transplanted mice as compared to the animals that received HSC gene therapy IV. This could be also due to competition between the ICV (transduced) transplanted cells and the un-transduced cells administered IV as rescue from conditioning, or to a pathological CNS milieu disfavoring the engraftment of the ICV transplanted HSPCs. Even if we did not test these hypotheses in our study, the evidence of ongoing inflammation and storage accumulation in the brain and spinal cord at time of transplant supports this view. Moreover, the biodistribution of ICV-transplanted HSPCs in the CLN1 disease mouse brain could be inefficient or uneven in some regions, which could explain the striking accumulation of AF storage in the cerebellum and in the thalamus. Similarly, limited engraftment in the spinal cord could be of relevance, as spinal cord pathology is considered of increasing importance in the Ppt1^-/-^ mouse^16^.

Based on these observations, we then decided to explore the therapeutic potential of combined (ICV and IV) administration of hPPT1 LV HSPCs. Importantly, the combined delivery of the transduced cells ICV and IV prevented disease manifestations and determined a long-lasting and thorough prevention of symptoms, a homogenous and substantial prevention of storage accumulation, and a widespread and broad prevention of cortical thinning, unlike that observed with the ICV-only approach and similarly to the IV-only group. Notably, neuroinflammation, measured by signal intensity of the CD68 marker, was more substantially reduced in ICV+IV treated mice compared to IV-or ICV-only recipients, suggesting a synergistic neuroprotective effect of the combined transplantation modality. These findings and the additive benefit of the combined ICV+IV HSPC gene therapy approach were supported by the PCAs performed on all histological as well as all behavioral analyses. Indeed, the ICV+IV group displayed histological and behavioral features more similar to WT animals than the other treatment modalities and a statistically superior therapeutic potential as compared to the IV-and ICV-only approaches.

Finally, gene-therapy was tested in late pre-symptomatic (130±15 days days old) animals to assess whether it could provide some benefit in such a challenging setting. Interestingly, the gene therapy retained therapeutic efficacy, despite, as foreseeably, the overall treatment outcome was less impactful than in young-adult mice. Indeed, the IV and ICV+IV transduced cell transplantation in symptomatic mice resulted in high donor-cell chimerism and Ppt1 reconstitution in the brain and spinal cord, similar to young adult mice. The disease progressed slowly and with milder symptoms (substantial absence of hindlimbs stiffness) in symptomatic-treated mice compared to UT controls, resulting in significantly increased overall survival of the former compared to the latter. Notably, IV+ICV treated mice significantly outlived not only the UT and mock-transplanted control mice, but also the IV treated cohort, further supporting the increased therapeutic potential of the former approach. Thus, in this challenging setting of aged, symptomatic mice treatment, despite not being likely able to rescue already established CNS damage, HSC gene therapy, and in particular the newly developed IV+ICV HSPC gene therapy approach, could prolong survival and delay disease manifestations of the affected mice, a remarkable result if compared to previously tested approaches that had resulted in therapeutic benefit only upon application to Ppt1^-/-^ newborns^22,32^, and in line with the results obtained with AAV gene therapy upon intrathecal administration, a route that allows widespread reconstitution of Ppt1 activity along the neuraxis^16^.

As far as safety is concerned, busulfan conditioning was feasible in CLN1 disease mice (despite the well-known pro-epileptogenic potential of this drug)^36^, provided that anti-seizure prophylaxis (e.g. benzodiazepine) is administered before and during the conditioning. Necropsy, hemocytometric and histopathological assessments performed on the animals from all treated groups did not report any findings associated with the transplantation of transduced HSPCs; only few animals (spread across all transplanted groups) had to be euthanized due to poor body conditions associated with the development of ascites and sarcomas in the abdomen. This was likely due to the well known long-term effects of busulfan (administered intraperitoneally diluted in a long-lasting solvent -see methods) described also in other long-term studies.

In conclusion, our study provides first evidence that: i) transplantation of wild type HSPCs exerts a partial but long-lasting mitigation of the symptoms and could thus represent *per se* a valuable approach for CLN1 disease if applied in pre-symtomatic stage and upon a proper conditioning regimen; ii) highly efficient gene transfer into the transplanted HSPCs enhances therapeutic benefit as compared to wild type cell transplant, in CLN1 disease as in other LSDs, with first demonstration of this dose-effect benefit for a purely neurodegenerative condition; iii) transplantation of hPPT1 over-expressing HSPCs by the novel ICV approach is sufficient to transiently ameliorate CLN1 disease symptomatology in the absence of hematopoietic tissue engraftment of the transduced cells; iv) the combinatorial transplantation of transduced HSPCs intravenously and ICV results in the most robust therapeutic benefit among the tested approaches on both pre-symptomatic as well as symptomatic animals, since it resulted in complete abrogation of the disease in a clinically relevant mouse model and determined a long-lasting and thorough prevention of symptoms. Notably, this same approach could uniquely benefit adult symptomatic animals, a finding of outmost importance for translation purposes. The clinical translatability of our strategy is further supported by the favorable safety profile we here showed.

## Materials and methods

### Lentiviral vector expressing hPPT1 gene

Human codon-optimized PPT1 was synthesized by Genewiz and cloned into a pCCLsin.cPPT.humanPGK.Wpre vector by BamHI/SalI digestion. Lentiviral vectors were produced and titered according to previous published protocols^20^.

### In vivo experiments

Experiments were performed on B6.129S6-Ppt1tm1Hof/SopJ mice (heareafter called Ppt1^-/-^), wild type C57BL/6J mice or B6.SJL-Ptprca Pepcb/BoyJ (hereafter called CD45.1 C57) mice, obtained from Jackson Lab and maintained at San Raffaele Hospital or at the Boston Children’s Hospital animal research facility.

### Isolation and transduction of murine hematopoietic stem cells

5-8 weeks old Ppt1^-/-^, wild type C57BL/6J or CD45.1 C57 donor mice were euthanized with CO_2_, and BM was harvested by crushing the femurs, tibias, humerus and iliac crest. HSPCs were purified by Lineage-(Lin-) selection using the Miltenyi Biotec Lineage Cell Depletion Kit with Magnetic separation with the autoMACS™ Separator, following manufacturer’s instruction. Isolated Lin-were transduced using different Lentiviral Vectors (LVs), for 16 hours at Multiplicity of Infection (MOI) 100 in Stem Span culture medium with antibiotics and cytokines (IL-3, IL-6, FLT-3 and SCF) at 37° C. The following LVs were used: pCCLsin.cPPT.humanPGK.DNGFR.Wpre (DNGFR-LV) for Lin-retrieved from wild type C57BL/6J mice; pCCLsin.cPPT.humanPGK.human-codon-optimized-PPT1.Wpre (hPPT1-LV) for Lin-retrieved from Ppt1^-/-^ mice.

A fraction of the transduced cells was used for colony forming assay to confirm the clonogenic potential as described^20^ whereas another fraction of transduced cells was cultured for 14 days ^20^ in order to assess vector copy number by digital qPCR and transgene expression by flow cytometry (in the case of DNGFR-LV) or through PPT1 enzymatic activity assay (for hPPT1-LV).

### Transplantation of murine hematopoietic stem cells

Six-eight weeks old Ppt1^-/-^ or wild type C57BL/6J mice were used as recipients of HSPCs transplantation. Before transplantation, mice were pre-treated with a myeloablative busulfan dose (27 mg/kg i.p. for 4 consecutive days) in order to ensure a sustained CNS engraftment of the transduced cells. Seizures are well-recognized complications of high-dose busulfan therapy in clinical practice. Given the intrinsic higher susceptibility to seizures observed in CLN1 patients and recapitulated in Ppt1^-/-^ mice, we applied seizure prophylaxis by benzodiazepine in Ppt1^-/-^ animals undergoing busulfan administration as strategy to reduce morbidity pre-transplant. The drug (diazepam) and administration protocol were defined based on literature evidence as follows: diazepam was administered for 10 days in the drinking water, starting two days before the first busulfan administration. 0.25 mg/kg diazepam was administered for 8 days, followed by two days of wash-out with 50% consecutive reductions in the administered dose (0.125 mg/kg and 0.063mg/kg). Lin-cells were injected IV (0.8 -1.2×10^6^ cells/mouse) or ICV (0.2 - 0.4×10^6^ cells/mouse) 24h after the last busulfan administration. To ensure proper reconstitution of the hematopoietic system in animals undergoing busulfan conditioning and receiving only the ICV administration of HSPCs, a systemic IV administration of freshly isolated untransduced BM cells after lysis (0.8 - 2.2×10^6^ cells/mouse), hereafter defined as “support BM cells” (retrieved from Ppt1 -/- mice or from CD45.1 mice), was performed 5 days after ICV transplantation.

ICV administration was performed by surgery; briefly: animals were anesthetized by isoflorane and mounted on a stereotaxic apparatus. Lin-cells (5 μl/injection site) were injected monolaterally in the lateral ventricles with a 27G Hamilton Syringe. Stereotaxic coordinates, referred to Bregma, were AP: 0.5, L:-1.0; DV: 2.5. After surgery, skin was sutured, and animals returned to their cages for recovery. After transplant, mice were monitored with collection of clinical signs for early identification of intercurrent deaths (ICDs) 3 days/week. In the case of ICDs, proper assessments were conducted as described below. Inter-current deaths up to day 31 are expected as consequence of the conditioning regimen, therefore no specific analyses were performed for ICDs before 31 days post-transplant (body was kept in formalin). ICDs or mice sacrificed due to poor clinical conditions (> 15% weight loss and hunched posture) occurring from Day 31 up to the end of study were processed, whenever feasible, in order to analyze the following parameters: i) transduced and mock-transduced cell engraftment on bone marrow cells, any lesion and in brain and spinal cord samples; ii) reconstitution of PPT1 activity on PBMCs or bone marrow cells, brain and spinal cord samples; iii) necropsy, histopathological and immunohistochemical examinations on a selected set of tissues. Moreover, dedicated assessments were scheduled at precise time intervals post-transplant:

#### i) dose-proof at 5-7 weeks after transplantation

To assess proper hematopoietic reconstitution by the transplanted cell progeny, the following parameters were assessed: i) transduced cell engraftment (VCN on bulk colonies grown from peripheral blood); ii) hematological parameters (complete blood count) on peripheral blood.

#### ii) behavioral assessment starting from 12 weeks after transplantation up to study termination

In order to monitor the effect of hematopoietic stem cell transplantation and the gene therapy treatment on disease progression and survival, behavioral disease severity score assessment was performed weekly and animals were tested for rotarod performance once every four weeks. All behavioral analyses were performed in a blind manner.

### Behavioral analyses

Disease severity score (DSS) was assigned by a trained operator in blind. Several behavioral symptoms were summarized with a score that was proportional to the extent of disease severity (DSS ranging from zero, i.e. no symptoms, to 9, i.e. animal is in poor body conditions). The total score was obtained by summing up one point for each of the following symptoms detected: skin lesions or scratches, tail flick (abnormal tail posture), hind-limb or fore-limb muscle atrophy, abnormal hind-limb or fore-limb displacement when the animal is raised by the tail for maximum 15 sec, hind-paws or fore-paws clasping behavior, limbs stiffness.

Motor performance was assessed on constant speed rotarod. Animals were placed on a rotating bar (4rpm, constant speed) for up to maximum 60 sec. Time spent on the rotating bar was recorded in seconds. Animals falling from the rod before 60 sec were returned to their cage for 5min before repeating the test for a maximum of three trials. The best performance was recorded for subsequent analysis.

### Mouse tissue collection and processing for flow cytometry and histology

Mice were euthanized under deep anesthesia by extensive intra-cardiac perfusion with cold PBS for 10 minutes after clumping the femur. Organs were then collected and differentially processed. Bone marrow (BM) cells were collected from the clumped femur as described. Brain and spinal cord were removed and divided in two longitudinal halves. For immunofluorescence and immunohistochemistry analysis, one half was fixed for 24 hours in 4% PFA, embedded in OCT compound and stored at -80°C, after equilibration in sucrose gradients (from 10 to 30%). One 1 mm thick section of the other half of each tissue was fixed in formalin for histopathology. For flow cytometry analysis, cells from the not-fixed tissue were mechanically disaggregated to obtain a single cell suspension in 20ml of GKN/BSA buffer (8g/L NaCl, 0.4 g/L KCl, 1.42 g/L NaH2P04, 0.93 g/L Na2HP04, 2 g/L D+ Glucose, pH 7.4 + 0.002% BSA). Additionally, spleen, thymus, liver, any lesion, sciatic nerves and hindlimb muscles were carefully dissected and fixed in formalin for histopathology. The rest of the body was preserved in formalin for further investigations.

### Flow-cytometric analysis

Cells from BM and brain were analyzed by flow cytometry upon re-suspension in blocking solution (PBS 5%FBS, 1%BSA) and labeling at 4°C for 15 minutes with the following specific antibodies: rat APC.Cy7 anti-mouse CD45 (BD Pharmingen) 1:100; rat APC anti-mouse CD11b (eBiosciences) 1:150; rat PE-Cy7 anti-mouse Ly-6C (BD Bioscence) 1:150; rat PE-Cy7 anti-mouse MHC.cl.II (BD Bioscence) 1:150; mouse Alexa647 anti-human human CD271 (NGF Receptor) (BD Pharmingen). For characterization of lin-purity, the following antibodies were used: rat APC anti-mouse CD11b (eBiosciences) 1:100; rat APC anti-mouse Gr-1 (BD Bioscence) 1:100; rat APC anti-mouse Ter119 (BD Bioscence) 1:100; rat APC anti-mouse CD3 (BD Bioscence) 1:100; rat APC anti-mouse B220 (BD Bioscence) 1:100; rat PE anti-mouse Sca-1 (BD Bioscence) 1:100; rat PE-Cy7 anti-mouse c-kit (BD Bioscence) 1:100. For the exclusion of death cells, we used either 7-AAD (1mg/ml) or DAPI (10mg/ml) (Sigma-Aldrich). Cells were analyzed by LSR Fortessa (Beckton Dickinson).

### Immunofluoresce and immunohistochemistry

Brains were serially cut in the sagittal plane on a cryostat in 14μm sections; spinal cords were cut in the coronal plane in 20μm sections. Tissue slides were washed twice with PBS, then sections were processed as follows:

#### CD271 (NGFR) staining

Sections were blocked with 0.3% Triton, 2% BSA, 10% NGS (Vector Laboratories) for 1 hour, then incubated O/N at 4C in blocking buffer containing the mouse anti-human CD271 (BD Pharmingen) biotinylated primary antibody. Afterwards, sections were washed trice with PBS and incubated for 30 min at RT with streptavidin-HRP (Perkin Elmer) dil. 1:100, washed and finally incubated for 8 min at RT with Tyramide-Cy5 (Perkin Elmer) dil. 1:300 in Amplification diluent (Perkin Elmer).

#### CD68 staining

Firstly, endogenous peroxydases were quenched by incubating the sections in 1% H_2_O_2_ in PBS for 10min. Sections were blocked with 0.3% Triton, 10% FBS (Vector Laboratories) for 1 hour, then incubated O/N at 4C in blocking buffer containing the rat anti-mouse CD68 (Abd Serotec) dil. 1:200, washed and incubated for 1h at RT with biotinylated anti-rat secondary antibody (Vector) dil. 1:200 in 1% FBS, in PBS. Afterwards, sections were washed trice with PBS and incubated for 1h at RT with Avidin-Biotin complex (Vector) and then with DAB/H_2_O_2_ mix (Sigma-Aldrich).

#### Neurotrace fluorescent NISSL staining

Sections were incubated for 10 min at RT in Neurotrace 660 (Molecular Probes, Invitrogen) dil. 1:250 in PBS, then washed trice with PBS.

Nuclei were stained with DAPI (Sigma) 0.5 mg/ml in PBS. Slices were washed in PBS, air dried and mounted with Mowiol.

#### NISSL staning

was performed as previously described^21^.

### Image acquisition and analysis

Sections stained by immunohistochemistry were acquired at a Nikon inverted microscope whereas immunofluorescne was acquired at a epifluorescence Nikon inverted microscope or a SP5 Leica confocal microscope equipped with laser lines: Ar-Kr (488 nm), He-Ne green (532 nm) and a UV diode.

Cortical thickness was quantified on 10x tilescan pictures of the brain (at least two slices per sample) with Fiji software. Briefly, the length of eleven segments drawn perpendicular to the cortical meninges and evenly spaced throughout the full cortical surface (from visual, to somatosensory and somatomotor cortex) was measured. The average of 3-4 segments per region was used in the analyses.

Autofluorescence storage material was quantified on 10x epifluorescence images acquired in the FITC channel in the cortex, hippocampus, thalamus and cerebellum of each sample (at least two slices per animal, four images per brain region). Background was subtracted by Fiji software, then CellProfiler was used to identify the autofluorescence positive signal.

CD68+ signal was quantified by CellProfiler on 10x brightfield images taken from the cortex and hippocampus of each animal (at least two slices/mouse). Fiji and Cellprofiler analysis pipelines are available on demand.

### Ppt1 enzymatic activity assay

Enzymatic activity was measured according to published protocols^10^ on proteins extracted by 6 freeze-thaw cycles in saline^20^ from cell pellets obtained from PB, BM, lin-liquid cultures, or through homogenization by sonication^22^ from brain and spinal cord tissue. The activity was normalized to the total protein content of the specimen.

### Statistical analysis

All statistical tests were two-sided. Normality of the samples was first verified by applying Shapiro-Wilk test. Then, Student’s t test or Mann-Whitney non-parametric test was used for two-group comparisons. For comparisons with more than two groups, one-way ANOVA with Tukey post hoc test (or Kruskal-Wallis non-parametric test followed by multiple-comparison corrected Dunn’s test) was used; for multiple variable comparisons, two-way ANOVA was applied. In case of longitudinal behavioral data, repeated measures ANOVA was applied. Survival and end-points analysis were performed by Log-Rank test. Differences were considered statistically significant at a value of *p < 0.05, **p < 0.01, ***p < 0.001. In all figures with error bars, the graphs depict means ± SEM.

### Study approval

Procedures involving animals and their care were conducted in conformity with the institutional guidelines according to the international laws and policies (EEC Council Directive 86/609, OJ L 358, 1 Dec.12, 1987; NIH Guide for the Care and use of Laboratory Animals, U.S. National Research Council, 1996). The specific protocols covering the studies described in this paper were approved by the Italian Ministry of Health, an internal ethical committee at San Raffaele Hospital and the Boston Children’s Hospital Institutional Animal Care and Use Committee.

## Abbreviations

AF: autofluorescent storage
BM: bone marrow
DSS: disease severity score
HEP: humane end point
HSC: hematopoietic stem cell
HSPC: hematopoietic and progenitor stem cell
ICDs: intercurrent deaths
ICV: intra-cerebroventricular
INCL: infantile neuronal ceroid lipofuscinosis
IV: intravenous
Lin-: lineage negative
LSD: lysosomal storage disorder
MOI: multiplicity of infection
NCL: neuronal ceroid lipofuscinosis
PBMCs: peripheral blood mononuclear cells
PPT1: plmytoil protein thioesterase
UT: untreated
WT: wild type.

## Data availability

The data that support the findings of this study are available from the corresponding author, upon reasonable request.

## Acknowledgements

We acknowledge FRACTAL, Flow cytometry Resource (Milan), DFCI Flow Cytometry Core, Alembic of San Raffaele Hospital and DFCI Confocal Light Microscopy Cores, and the Animal Behavior & Physiology Core of Boston Children’s Hospital (in the person of Nick Andrews) for technical support. We wish to thank Danilo Pellin for assistance with statistical analyses.

## Author contributions

M.P. performed the experiments, performed the analyses, wrote the manuscript, drafted the figures; J.P., O.J.C., T.D.M., A.Z., R.M., E.C., S.D. managed the mouse colony, helped with *in vivo* behavioral assessments, enzymatic activity assays and VCN evaluations; R.K., J.P. and

T.D.M. helped with immunohistochemistry experiments; V.P. helped with histopathological studies and data interpretation; A.B. conceived the study, retrieved the funding, reviewed the manuscript.

## Funding

This work was supported by the following grants: ERC-2013-CoG, Project No 617162, and NIH R01 HD095935-04 to AB.

## Conflict of Interests

AB and MP are co-authors in the patent PCT/US2017/056774. The Authors have no other competing interests to declare.

## Figure legends

**Suppl. Fig. S1**.

**A**. Representative pictures of the disease manifestations displayed by symptomatic Ppt1^-/-^ mice during tail suspension. **B**. Progressive deterioration of animals body condition demonstrated by the DSS. * = p< 0.05; ** = p< 0.01; *** = p< 0.001 Repeated Measures ANOVA followed by Tukey’s post-hoc test. **C**. Progressive motor deficits in Ppt1^-/-^ mice assessed by rotarod test. **** = p < 0.0001 Repeated Measures ANOVA followed by Tukey’s post-hoc test. **D**. Comparison of DSS with different disease manifestations (limbs clasping, stiffness and rotarod deficits) and with survival by Log-rank analysis. See supplementary table 2 for statistics.

**Suppl. Fig. S2**.

**A**. Schematic representation of the transplant procedure and animals monitoring post-transplant. **B**. Longitudinal assessment of DSS in HSPCs or BM transplanted mice in comparison with mock transplanted or untreated Ppt1^-/-^ mice. **C**. Assessment of donor-cell engraftment in peripheral blood (PB) and BM of transplanted mice. **D**. Assessment of donor-cell engraftment in the brain of transplanted mice. **E**. Ppt1 enzymatic activity in brain samples from transplanted mice. **F**. Autofluorescent (AF) storage material in the brain of transplanted mice. **G**. Assessment of cortical thickness in transplanted mice.

**Suppl. Fig. S3**.

Representative brightfield microscope microphotographs of CD68 immunoreactive DAB staining in the cortex, hippocampus, thalamus and cerebellum, in Ppt1 ^+/+^ mice analyzed at 400 days of age, or Ppt1 ^-/-^ mice left untreated (UT) or mock transplanted with Ppt1 ^-/-^ unmanipulated HSPCs (analyzed at humane end point, ∼ 250 days of age); Ppt1 ^-/-^ mice transplanted with hPPT1-LV transduced HSPCs administered IV, ICV or ICV+IV, analyzed at study termination, 360-400 days of age. Scale bar = 200 µ m.

**Suppl. Fig. S4**.

**A**. VCN and hPPT1 activity retrieved in the BM of Ppt1 ^-/-^ transplanted with hPPT1-LV transduced HSPCs IV, ICV or ICV+IV, analyzed at study termination. **B**. Classification of intercurrent deaths (ICD) reported in Ppt1 ^-/-^ mice mock transplanted, transplanted with hPPT1-LV transduced HSPCs IV, ICV or ICV+IV, or untreated (UT). HEP = humane end point; FD = found dead; EOS = end of study. **C**. Hemocytometric analysis of the peripheral blood of transplanted or untreated Ppt1 ^-/-^ and Ppt1 ^+/+^ mice at 5-6 weeks post-transplant. **D**. Hemocytometric analysis of the peripheral blood of transplanted or untreated Ppt1 ^-/-^ and Ppt1 ^+/+^ mice at euthanasia. WBC = white blood cells; RBC = red blood cells; HGB = hemoglobin; HCT = hematocrit; MCV = mean corpuscular volume; MCH = mean corpuscular hemoglobin; MCHC = mean corpuscular hemoglobin concentration; PLT = platelets count.

## Notes

### Competing Interest Statement

Alessandra Biffi and Marco Peviani are co-authors in the patent PCT/US2017/056774. The Authors have no other competing interests to declare.

