## Supplemental Figures and tables for "An innovative hematopoietic stem cell gene therapy approach benefits CLN1 disease in the mouse model": Peviani et al 2022-02-28 Supplementary tables rev.docx

**Supplementary table 1. Example of DSS assessment.**

| **Symptoms** | **Present** | **Score** | **TOTAL** |
| --- | --- | --- | --- |
| Skin wounds | ✔ | +1 | 6 |
| Seizures (jerks, pop-corn) | ✔ | +1 |  |
| Hind-limbs abduction deficits (close to midline) | ✔ | +1 |  |
| Hind-limbs orizontal displacement |  |  |  |
| Hind-limbs clasping | ✔ | +1 |  |
| Hind-limbs muscle atrophy | ✔ | +1 |  |
| Fore-limbs muscle atrophy |  |  |  |
| Fore-limbs clasping |  |  |  |
| Limbs stiffness |  |  |  |
| Tail-flick | ✔ | +1 |  |

**Supplementary table 2. Log-rank analysis of DSS and disease symptoms manifestation.**

|  | **Fig. S1D - Log-rank analysis of disease symptoms** | | | | | | |
| --- | --- | --- | --- | --- | --- | --- | --- |
|  | **DSS ≥ 2** | **DSS ≥ 4** | **DSS ≥ 6** | **clasping** | **rotarod** | **stiffness** | **survival** |
| **DSS ≥ 2** |  |  |  |  |  |  |  |
| **DSS ≥ 4** | < 0.0001 |  |  |  |  |  |  |
| **DSS ≥ 6** | < 0.0001 | < 0.0001 |  |  |  |  |  |
| **clasping** | ns | < 0.0001 | < 0.0001 |  |  |  |  |
| **rotarod** | 0.0006 | ns | < 0.0001 | 0.0316 |  |  |  |
| **stiffness** | < 0.0001 | < 0.0001 | ns | < 0.0001 | < 0.0001 |  |  |
| **survival** | < 0.0001 | < 0.0001 | < 0.0001 | < 0.0001 | < 0.0001 | 0.0164 |  |

**Supplementary table 3. Statistics for Log-rank analyses among HSC gene therapy groups.**

|  | **Fig. 2C - Survival** | | | | |
| --- | --- | --- | --- | --- | --- |
|  | **IV tx** | **ICV tx** | **IV+ICV tx** | **Mock tx** | **Ppt1 -/- UT** |
| **IV tx** |  |  |  |  |  |
| **ICV tx** | ns |  |  |  |  |
| **IV+ICV tx** | ns | ns |  |  |  |
| **Mock tx** | <0.0001 | <0.0001 | <0.0001 |  |  |
| **Ppt1 -/- UT** | <0.0001 | <0.0001 | <0.0001 | ns |  |
|  | **Fig. 2E - DSS ≥ 4** | | | | |
|  | **IV tx** | **ICV tx** | **IV+ICV tx** | **Mock tx** | **Ppt1 -/- UT** |
| **IV tx** |  |  |  |  |  |
| **ICV tx** | 0.0034 |  |  |  |  |
| **IV+ICV tx** | ns | <0.0001 |  |  |  |
| **Mock tx** | 0.0009 | 0.0187 | 0.005 |  |  |
| **Ppt1 -/- UT** | <0.0001 | <0.0001 | <0.0001 | 0.0019 |  |
|  | **Fig. 2F - Limb stiffness** | | | | |
|  | **IV tx** | **ICV tx** | **IV+ICV tx** | **Mock tx** | **Ppt1 -/- UT** |
| **IV tx** |  |  |  |  |  |
| **ICV tx** | ns |  |  |  |  |
| **IV+ICV tx** | ns | ns |  |  |  |
| **Mock tx** | 0.0114 | 0.0107 | 0.0224 |  |  |
| **Ppt1 -/- UT** | 0.0073 | 0.0193 | 0.0129 | ns |  |
|  | **Fig. 5A - Survival** | | | | |
|  | **IV tx** | **IV+ICV tx** | **Mock tx** | **Ppt1 -/- UT** |  |
| **IV tx** |  |  |  |  |  |
| **IV+ICV tx** | 0.0066 |  |  |  |  |
| **Mock tx** | 0.0004 | <0.0001 |  |  |  |
| **Ppt1 -/- UT** | 0.0016 | <0.0001 | ns |  |  |
|  | **Fig. 5C – DSS ≥ 4** | | | | |
|  | **IV tx** | **IV+ICV tx** | **Mock tx** | **Ppt1 -/- UT** |  |
| **IV tx** |  |  |  |  |  |
| **IV+ICV tx** | ns |  |  |  |  |
| **Mock tx** | ns | ns |  |  |  |
| **Ppt1 -/- UT** | ns | ns | ns |  |  |
|  | **Fig. 5D – Limbs stiffness** | | | | |
|  | **IV tx** | **IV+ICV tx** | **Mock tx** | **Ppt1 -/- UT** |  |
| **IV tx** |  |  |  |  |  |
| **IV+ICV tx** | ns |  |  |  |  |
| **Mock tx** | 0.0048 | <0.0001 |  |  |  |
| **Ppt1 -/- UT** | 0.0002 | <0.0001 | ns |  |  |

**Supplementary table 4. Macroscopic histopathological findings in the abdominal cavity in mice .**

| Group name | IV tx | ICV tx | IV+ICV tx | Mock tx | Ppt1 -/- UT | Ppt1 +/+ tx | Ppt1 +/+ UT |
| --- | --- | --- | --- | --- | --- | --- | --- |
| Treatment | Ppt1^-/-^ + HSCPs PPT1-LV (IV) | Ppt1^-/-^ + HSCPs PPT1-LV (ICV) | Ppt1^-/-^ + HSCPs PPT1-LV (IV+ICV) | Ppt1^-/-^ + autologous un-transduced HSCPs IV | Ppt1^-/-^ un-transplanted | Ppt1^-/-^ + autologous un-transduced HSCPs IV | Ppt1^+/+^ un-transplanted |
| Number of animals in group | 13 | 11 | 10 | 11 | 8 | 3 | 3 |
| Mass | 2 | 1 | 3 | 0 | 0 | 1 | 1 |

**Supplementary table 5. Microscopic histopathological findings in the abdominal cavity in mice .**

| Group name | IV tx | ICV tx | IV+ICV tx | Mock tx | Ppt1 -/- UT | Ppt1 +/+ tx | Ppt1 +/+ UT |
| --- | --- | --- | --- | --- | --- | --- | --- |
| Treatment | Ppt1^-/-^ + HSCPs PPT1-LV (IV) | Ppt1^-/-^ + HSCPs PPT1-LV (ICV) | Ppt1^-/-^ + HSCPs PPT1-LV (IV+ICV) | Ppt1^-/-^ + autologous un-transduced HSCPs IV | Ppt1^-/-^ un-transplanted | Ppt1^-/-^ + autologous un-transduced HSCPs IV | Ppt1^+/+^ un-transplanted |
| Number of animals in group | 13 | 11 | 10 | 11 | 8 | 3 | 3 |
| Sarcoma NOS | 3 | 1 | 3 | 0 | 0 | 1^a^ | 0 |
| Atypical Fibroplasia | 1 | 0 | 0 | 1 | 0 | 1^a^ | 0 |
| Total | 4 | 1 | 3 | 1 | 0 | 1 | 0 |

^a^ sarcoma and atypical fibroplasia occurred in the same animal
