## Supplementary figures and images for "An innovative hematopoietic stem cell gene therapy approach benefits CLN1 disease in the mouse model"

### Figure S1.tif

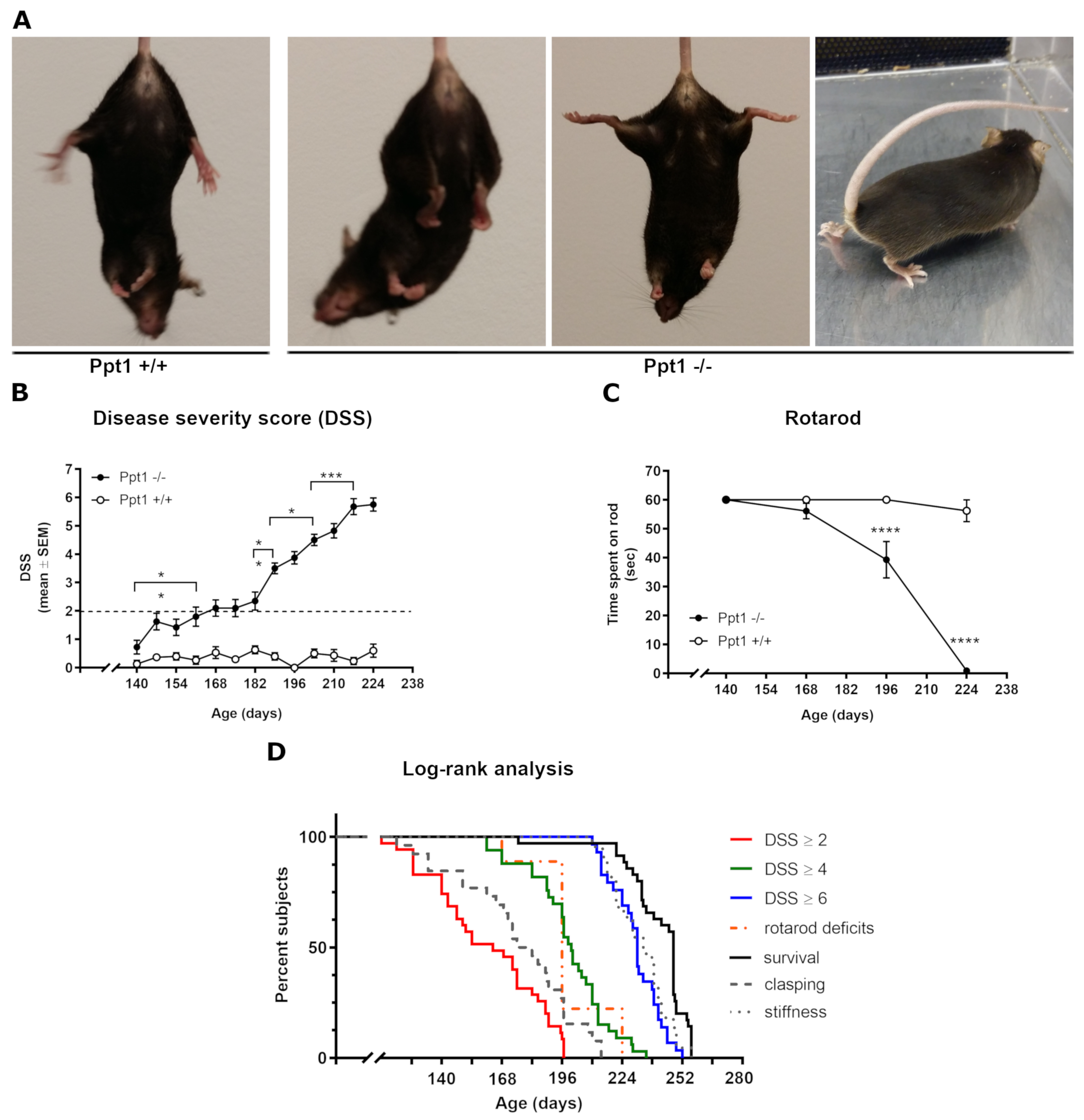

### Figure S2.tif

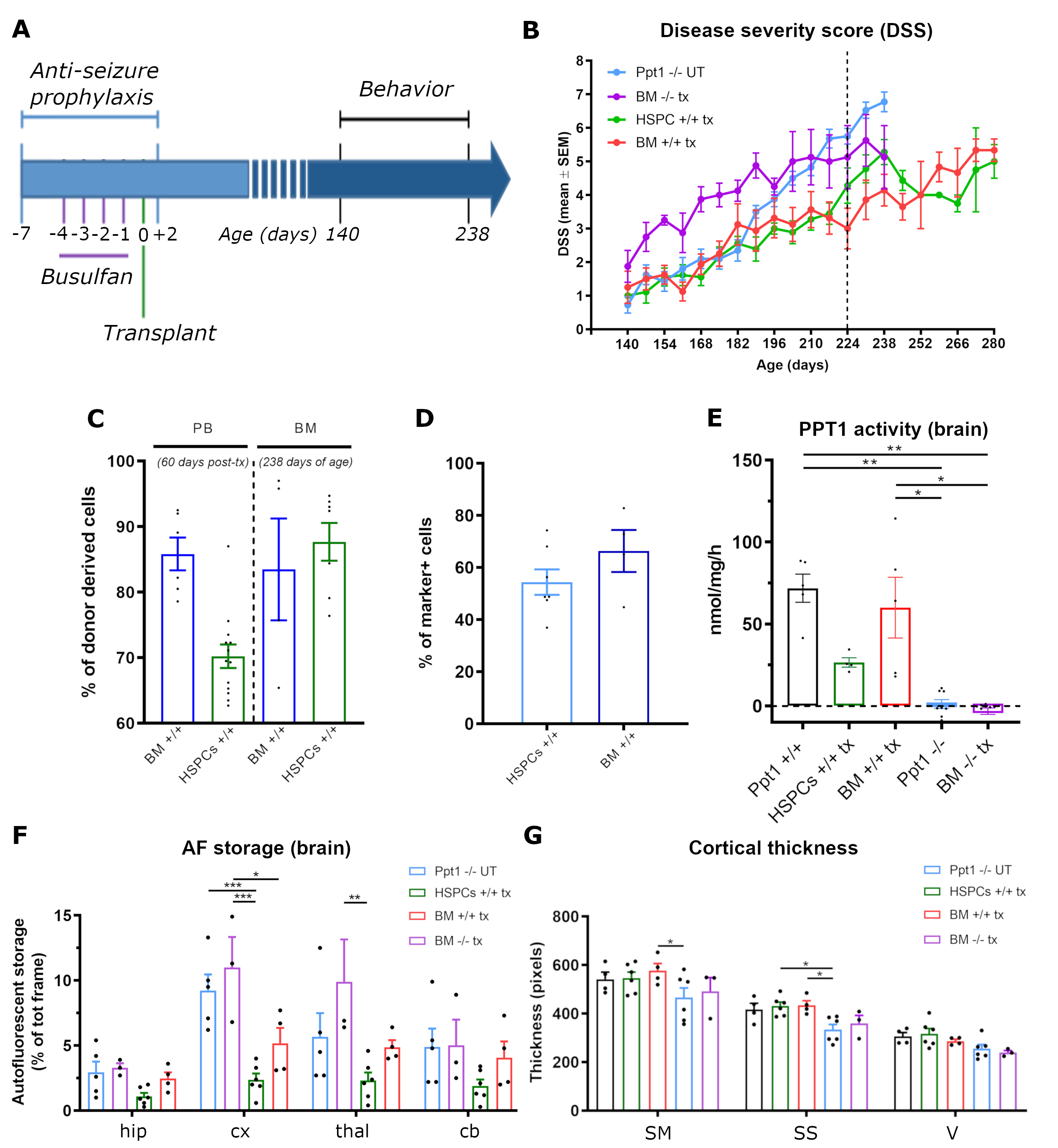

### Figure S3.tif

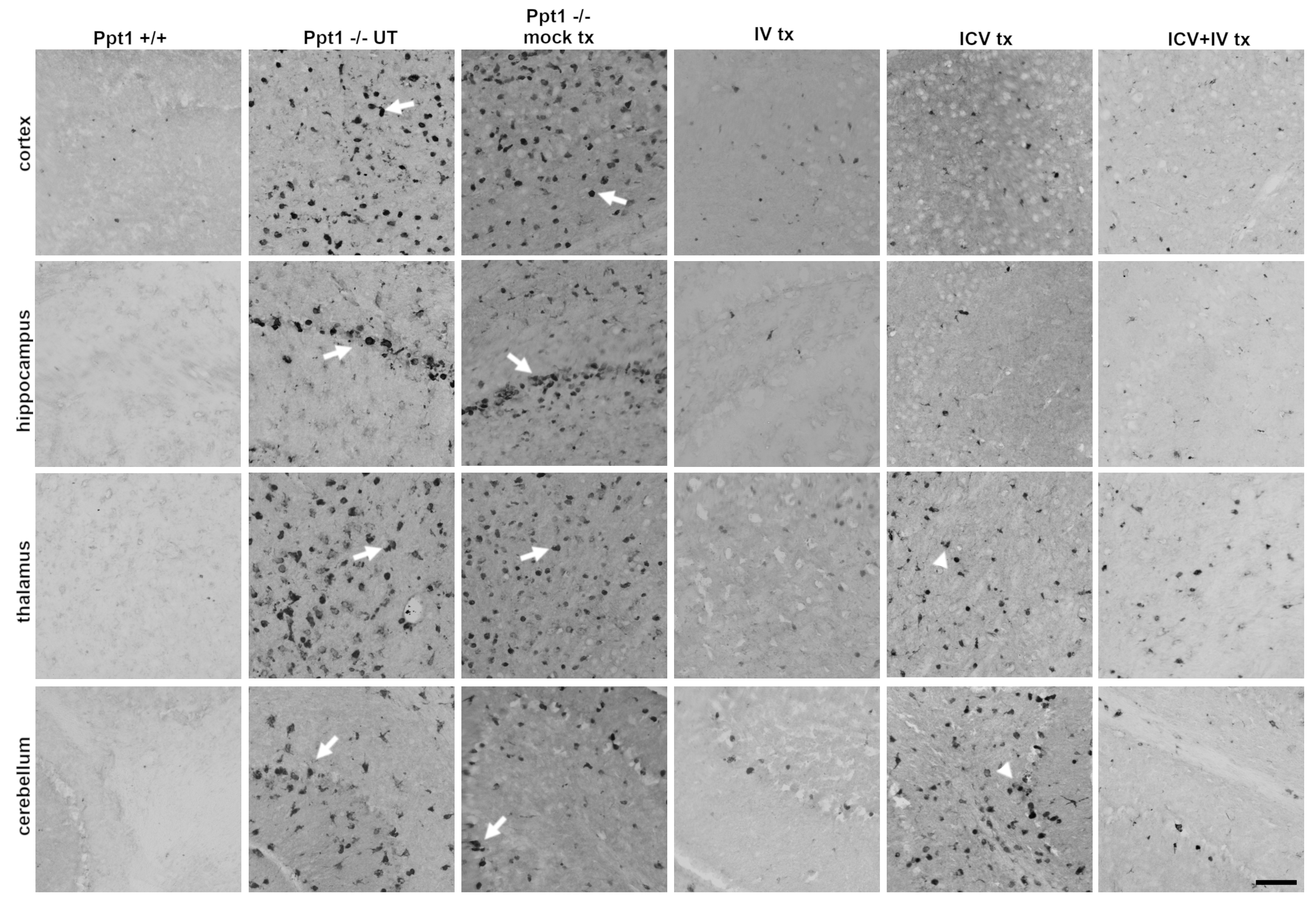

### Figure S4.tif

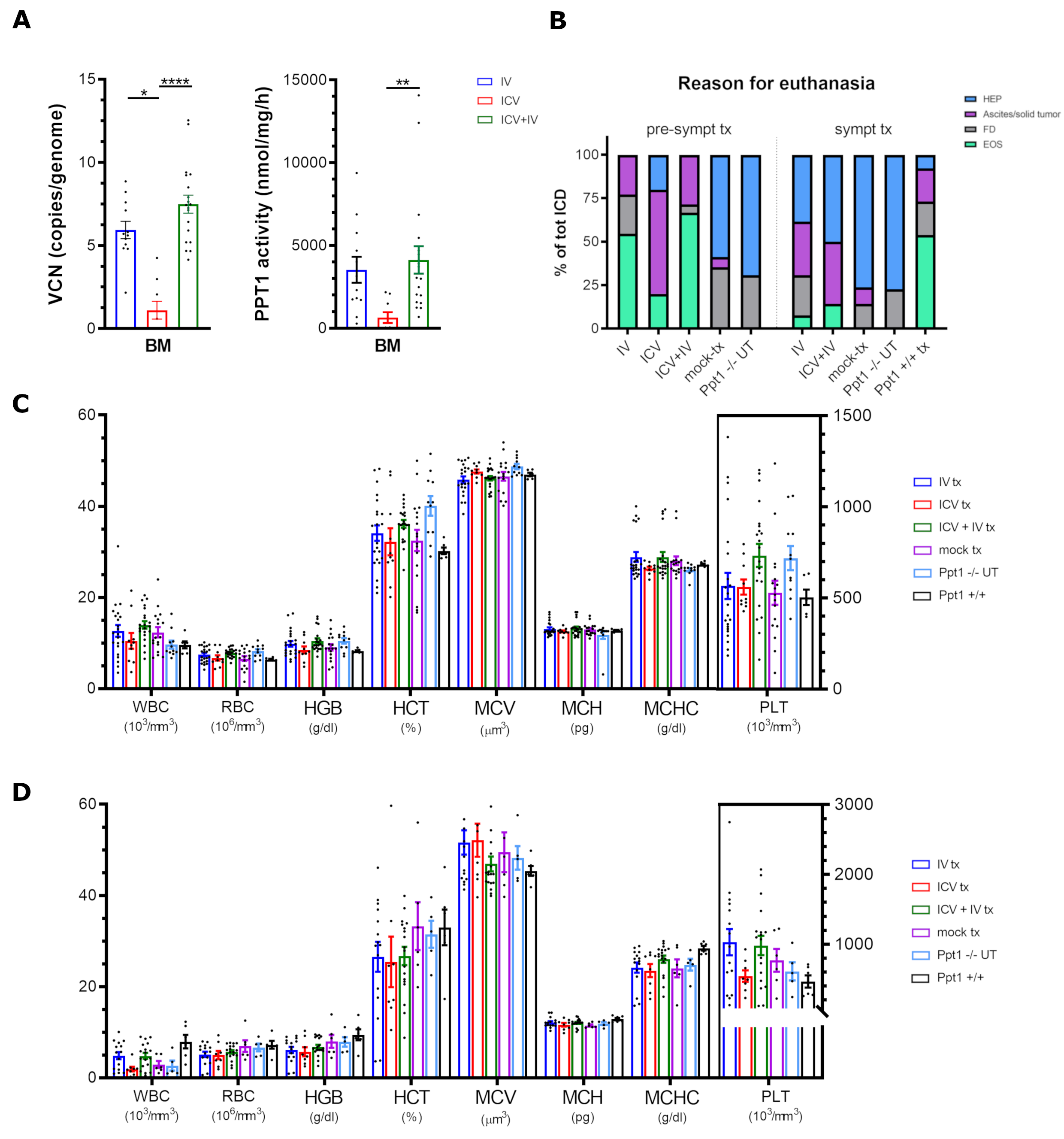
